# A novel high-throughput single-cell DNA sequencing method reveals hidden genomic heterogeneity in the unicellular eukaryote *Leishmania*

**DOI:** 10.1101/2025.09.10.675331

**Authors:** Gabriel H. Negreira, Pieter Monsieurs, Jean-Claude Dujardin, Malgorzata A. Domagalska

## Abstract

Genome instability is considered a major driver of adaptation in eukaryotic microorganisms, but its study is hampered by the limited availability of genomic technologies with single-cell resolution. Here we present a novel high throughput method to reconstruct both structural and nucleotide information at single-cell level in *Leishmania*, a protozoan parasite with a remarkable genome plasticity characterized by frequent gene copy number variations (CNVs) and high aneuploidy mosaicism. By combining the use of semi-permeable capsules with primary template-directed amplification, we could determine the karyotypes of hundreds of *Leishmania* parasites, detect distinct CNVs between different cell populations, and identify sub population of cells harboring distinct nucleotide variants, including in genes associated with drug resistance. This approach provides a powerful new framework to uncover hidden evolutionary potential in complex microbial populations, with application in studying adaptation, drug resistance, and genome evolution in *Leishmania* and other pathogens.

## INTRODUCTION

Genome instability plays a paradoxical and yet pivotal role in the evolutionary success of many unicellular pathogens. While genomic integrity is typically vital for cellular function, moderate levels of instability – either through intra-chromosomal copy number variations (CNV) or through dosage changes of whole chromosomes (aneuploidy) – can facilitate survival under environmental stress by quickly diversifying phenotypes in a cell population^1–3^. For instance, aneuploidy promotes survival to drug pressure and host immunity in *Candida albicans*^4,5^, and CNVs were linked to survival under nitrogen-limiting conditions in *Saccharomyces cerevisiae*^6^. Aneuploidy and/or CNVs are also commonly found in cancer and are linked to tumor progression and drug resistance ^7,8^. Moreover, these types of structural variation are also found in protist parasites such as *Giardia intestinalis*, *Plasmodium spp.*, *Trypanosoma cruzi*, and *Leishmania*^9–13^, representing potential adaptive mechanisms in clinically relevant organisms.

*Leishmania* is a protist parasite that cause leishmaniasis, a neglected tropical disease which affects more than 99 countries with 700,000-1,000,000 new cases every year^14^. These parasites also represent an important model to study genome instability in eukaryotes, owing to its remarkable genomic plasticity and the central role that genome instability plays in its biology. First, while most eukaryotes maintain a euploid genome, i.e., with equal copy numbers of each chromosome, *Leishmania* breaks this rule, as aneuploidy is not an exception but the standard for these parasites. In its most basic form, aneuploidy is present as a characteristic polysomy of chromosome 31 contrasting with an otherwise fully diploid genome^15^. However, high levels of chromosome instability lead to frequent dosage changes involving virtually any of the 36 chromosomes, making any *Leishmania* population a ‘mosaic’ of distinct aneuploid karyotypes^16,17^. Aneuploidy patterns can shift in response to environmental stresses ^18–21^ and are directly reflected into RNA levels^18,22^ and partially at protein levels^23^. In addition, *Leishmania* genome instability is also expressed in the form of frequent CNVs. Small segments of 20–70 kb can form extrachromosomal circular or linear amplicons carrying genes, including those encoding drug resistance factors^24^. Expansion of intra-chromosomal loci is also common. For instance, a common CNV found in chromosome 23 of some *L. donovani* strains was linked to resistance to antimonials in strains from Nepal^19,25^. In addition, the occurrence of mixed infections – where multiple *Leishmania* strains or even species coexist in the same host^26–28^ – further highlight the need for genomic approaches that resolve parasite genotypes at single-cell resolution.

In a previous study our group established the first single-cell DNA sequencing (scDNAseq) study with *Leishmania*^29^. Using single parasites isolated in plates with fluorescence activated cell sorting (FACS), our group could successfully sequence 47 parasites, of which 28 had their karyotypes determined. This study also demonstrated that the whole-genome amplification (WGA) method must be chosen based on the downstream analysis, with multiple displacement amplification (MDA) being more suited for nucleotide sequence information, and PicoPlex^TM^ performing better for structural variations such as chromosome copy number assessment and CNVs^29^. Later, we described the application of the Chromium single-cell CNV solution (10X Genomics^TM^) as a high throughput, emulsion droplet-based approach that could reconstruct karyotypes in thousands of *Leishmania* cells, revealing potential paths towards karyotype diversification, but with no access to nucleotide variant detection due to shallow genomic coverage ^17^. Moreover, the Chromium single-cell CNV solution was discontinued in 2020, with no replacement currently available to our knowledge. This gap highlights the urgent need for novel, high-throughput scDNAseq approaches.

Here we developed a novel high throughput scDNAseq method tailored to detecting structural and nucleotide variants at single-cell level in eukaryotic parasites such as *Leishmania*. Our method is based on the commercially available Single-Microbe DNA Barcoding solution (Atrandi Biosciences^TM^), which originally offers a workflow for cell encapsulation using semi-permeable capsules (SPCs), whole genome amplification (WGA) and genome barcoding of bacterial cells. The main advantage of SPCs over emulsion droplet capturing lies in its semi-permeable nature: large biomolecules (such as genomic DNA or whole organelles) are retained inside the capsule, while small molecules (e.g. enzymes, buffers, cofactors) can diffuse in and out^30^. Moreover, SPCs are resistant to harsh lysis chemicals, deep freezing and heat incubations. These properties permit multi-step reactions to be performed in the pools of encapsulated cells by simply washing or exchanging the surrounding solution, making custom protocol modifications easy. We compared the performance of the method with libraries previously made with the Chromium single-cell CNV, showing that the original SPC-based protocol successfully generates scDNAseq libraries with *Leishmania*, but its uneven amplification prevents determination of structural variants. We solved this issue by using primary template-directed amplification (PTA)^31^ as alternative WGA. This modification allowed us to fully reconstruct the karyotype of hundreds of *L. donovani* parasites and to detect distinct CNV patterns between cells. Moreover, the higher coverage seen in PTA unlocked the ability to simultaneously retrieve nucleotide sequence information from single-cells, which was not possible before. We demonstrate the ability of the method to distinguish parasite subpopulations of the same species, as well as small populations in the same strain, revealing minor genotypic variants among cells in clinical isolates, including variants in genes related to drug-resistance in small subpopulations of parasites. Together, our results represent a significant advancement in single-cell genomics for *Leishmania* and other eukaryotes, providing a scalable, high-resolution tool to dissect both structural and sequence-level variation, and to deepen our understanding of parasite heterogeneity, evolution, and adaptation.

## MATERIALS AND METHODS

### Parasite Culture

In the present study, we refer to terms such as population, strain and clone, according to the nomenclature proposed for salivarian trypanosomes^32^. Accordingly: (i) a population is a group of *Leishmania* cells present at a given time in a given environment (e.g., culture or host); (ii) a strain is a population derived by serial passage *in vitro* from a primary isolate (in this case, from clinical samples) without any implication of homogeneity but with some degree of characterization (here, bulk genome sequencing); (iii) a clone is a population of cells derived from a single individual presumably by binary fission. Here, two *L. donovani* strains isolated from visceral leishmaniasis patients were used, the BPK081/0 cl8 (MHOM/NP/02/BPK081/0) strain, which was isolated in Nepal, and HU3 (MHOM/ET/67), isolated in Ethiopia. Both strains were retained at the cryobank of the Institute of Tropical Medicine Antwerp and were maintained in HOMEM medium supplemented with 20% fetal bovine serum at 26 °C with periodic passages done every 7 days in 1 in 50 dilutions for downstream experiments. The BPK081/0 cl8 strain was submitted to single-cell sequencing 7 passages after cloning, while HU3 had an unknown of passages (minimum of 15 passages).

### Single-cell encapsulation into SPCs

Single-cell encapsulation was done using semi-permeable capsules (SPC). Promastigotes from each strain were collected independently at late-stationary phase (day 7) by centrifugation at 1500 rcf for 5 minutes followed by 3 washes with PBS (magnesium and calcium-free) and passed through a 5 µm strainer to remove cell clumps. Then, cell concentration was adjusted to 10^7^ parasites/ml in each strain, and both were combined at 50/50 ratio for downstream applications. The cell mixture was submitted to SPC encapsulation using the Flux device (Atrandi), following the Flux SPC Generator protocol (V6.3), with 6.3 µl of the cell suspension loaded, targeting a total of 15.000 encapsulated cells (corresponding to ∼5.000 sequenced cells). After encapsulation, SPCs were washed 3 times with 1 ml of Capsule Wash Buffer (CWB - Atrandi) and were stored at −80°C overnight. Afterwards, SPCs were thawed back to room temperature, washed 3 times with 1 ml of Wash Buffer (WB - Atrandi) and resuspended in 480 µl WB. The SPC suspension was subsequently divided into 6 samples with 80 µl each which were maintained on ice. Samples were named according to the WGA method applied: SPC-STD1-4 for the samples submitted to the standard WGA of the Single-Microbe Genome Barcoding Kit, and SPC-PTA1-2 for the samples submitted to PTA. A summary of lysis and WGA conditions is provided in the results section (Table 1).

**Table 1.** - Summary of the different protocol modifications and results of each sample. Lysis Std: Standard lysis protocol of the Single-Microbe DNA Barcoding kit, i.e, 0.8 M KOH lysis for 15 minutes at 23°C. Lysis 1h: Same as standard lysis but with 1h incubation instead of 15 minutes. Lysis 99 °C: same as standard lysis but with incubation at 99 °C instead of 23 °C. Lysis PTA: lysis protocol of the ResolveDNAv1 kit. WGA Std: Standard WGA of the Single- Microbe DNA Barcoding kit, which is based on MDA. Estimated doublets: doublets estimated by assuming random pairing assuming random cell pairing, dividing the observed frequency of HU3/BPK081 doublets by the expected proportion of total doublets, 2pq, where p and q are the relative frequencies of HU3 and BPK081 singlets respectively.

| sample | Methanol Fixation | Lysis | WGA | Total Reads | SPCs | true cells | Reads in true cells | Median reads per cell | HU3 cells | BPK081 cells | Detected doublets | Estimated doublets |
| --- | --- | --- | --- | --- | --- | --- | --- | --- | --- | --- | --- | --- |
| SPC-STD1 | yes | Std | Std | 33,050,073 | 184 | 147 | 31,548,327 | 188,688 | 79 | 62 | 6 | 12 |
| SPC-STD2 | no | Std | Std | 55,394,197 | 257 | 205 | 52,847,223 | 197,432 | 113 | 88 | 4 | 8 |
| SPC-STD3 | no | 1h | Std | 61,026,392 | 317 | 221 | 56,738,608 | 239,455 | 110 | 106 | 5 | 10 |
| SPC-STD4 | no | 99 °C | Std | 1,834,390 | 95 | 70 | 1,712,470 | 20,128 | 0 | 27 | 43 | NA |
| SPC-PTA1 | no | Std | PTA | 251,152,697 | 742 | 161 | 238,010,354 | 1,116,078 | 88 | 71 | 2 | 4 |
| SPC-PTA2 | no | PTA | PTA | 215,348,910 | 869 | 253 | 190,957,268 | 718,145 | 146 | 101 | 6 | 12 |

### Cell lysis

Of the samples described above, 5 were submitted to different variations of the lysis protocol described in the Single-Microbe DNA Barcoding user guide (Atrandi) version 4.1 (P/N: CKP-BARK1): sample SPC-STD1 was first fixed by slowly adding 900 µl of ice-cold methanol to the SPC suspension on ice, following by an incubation at −20 °C for 30 minutes, 3 washes with 1 ml WB, and a final resuspension in 80 µl of WB. For samples SPC-STD2-4 and SPC-PTA1-2, methanol fixation was skipped. Then, the 5 samples received 80 µl of 2X Lysis Buffer (0.8 M KOH, 20 mM EDTA, 200 mM DTT - Atrandi), with samples SPC-STD1, SPC-STD2, and SPC-PTA1 being incubated at 23 °C for 15 minutes, sample SPC-STD3 being incubated at 23 °C for 60 minutes, and sample SPC-STD4 being incubated at 99 °C for 15 minutes for lysis. After lysis, each sample was washed 3 times with 160 µl of Neutralization Buffer (NB - Atrandi) followed by 3 washes with WB, with final resuspension in 25 µl WB, with tubes being kept on ice.

For sample SPC-PTA2, the lysis protocol from the ResolveDNA kit (BioSkryb) was followed instead. In summary, SPCs were washed 3 times with 25 µl of Cell Buffer (CB – BioSkryb). Then, 25 µl of the MS mix (BioSkryb) were added, followed by an incubation at room temperature for 20 minutes at 1400 rpm in a thermomixer. Then, SPCs received 25 µl of SN1 reagent (BioSkryb) with another room temperature incubation for 1 minute at 1400 rpm followed the addition of 25 µl of the SDX reagent (BioSkryb) with a 10-minute incubation at room temperature without agitation. After lysis, SPCs were kept on ice until PTA. Since the components of the PTA lysis protocol could affect the downstream PTA, we also added the aforementioned lysis reagents from the PTA protocol to SPC-PTA1, skipping the incubation times.

### Whole-genome amplification

Samples SPC-STD1-4 were submitted to the standard WGA method described in the Single-Microbe DNA Barcoding user guide (Atrandi). In summary, the 25 µl SPC suspensions were combined with 16,25 µl of nuclease-free water, 6.25 µl of 10X WGA Reaction Buffer (Atrandi), 6.25 µl of dNTP Mix (Atrandi), 3.125 µl Primer mix (Atrandi), 0.625 µl of 0.1 M DTT, 0.625 µl of 10% Pluronic F-68, 1.25 µl of WGA Enhancer (Atrandi), and 3.125 µl of WGA Polymerase (Atrandi) on ice. The WGA mixture was then submitted to an incubation at 45 °C for 15 minutes followed by 65 °C for 10 minutes in a thermocycler and were returned to ice.

For samples SPC-PTA1-2, the PTA protocol from the ResolveDNA kit version 1 (BioSkryb) was followed instead. In summary, a PTA master mix was prepared on ice by combining 85 µl of SB4 reagent, 17 µl of 1X SS2 reagent, 13,6 µl of SEZ1 reagent, and 20,4 µl of SEZ2 reagent (all from ResolveDNA kit). Then, 66,7 µl of the PTA master mix was added to the 100 µl of lysed SPCs of samples SPC-PTA1-2. These samples were then submitted to a 10-hour incubation at 30 °C followed by an inactivation at 65 °C for 3 minutes.

After WGA, all samples were washed 3 times with 200 µl of WB, with final resuspension done in 25 µl. DNA amplification was evaluated in all samples by staining 1 µl of SPCs with 5µM SYTO9 (ThermoFisher) and visualizing SPC positives in a fluorescence microscope. For samples SPC-STD1-4, amplified DNA was debranched by combining the 25 µl of SPC suspension with 17.5 µl of Nuclease-free water, 5 µl of 10X Debranching Buffer (Atrandi) and 2.5 µl of Debranching Enzyme (Atrandi) followed by an incubation at 37 °C for 1 hour at 1000 rpm agitation in a thermomixer. After debranching, SPCs were again washed 3 times with WB with final volume of 25 µl. The samples submitted to PTA SPC-PTA1-2 did not require debranching and were directly submitted to barcoding instead.

### Barcoding of amplified genomes

Barcoding of amplified SPC-trapped DNA was done following the split-pool barcoding approach described in the Single-Microbe DNA Barcoding user guide (Atrandi) for all 6 samples. First, DNA end preparation was done to allow the subsequent ligation of barcodes to the amplified DNA molecules. This was done by adding 3.5 µl of End Prep Buffer (Atrandi), and 1.5 µl of End Prep Enzyme (Atrandi) to the 25 µl of SPC suspensions, followed by an incubation at 20 °C for 30 minutes and inactivation at 65 °C for 30 minutes. Each sample immediately received 25 µl of a ligation master mix (77 µl of nuclease-free water, 66 µl of 10X ligation buffer (Atrandi) and 22 µl of Ligation Enzyme (Atrandi) and was distributed along 4 wells (10 µl per well) of the first-round columns of the barcoding plate from the Single-Microbe DNA Barcoding kit (Atrandi). The well positions of each sample was recorded to be used as sample index later. Barcode ligation was carried out at 25 °C for 15 minutes and stopped with 40 µl of 1X Stop Buffer (SB - Atrandi) with a 5-minute incubation at room temperature. After that, SPCs from all wells were pooled in a single tube, washed 5 times with 1 ml WB (final volume of 150 µl), combined with 150 µl of a new ligation master mix, and redistributed along the 24 wells of the second-round columns in the barcoding plate (10 µl per well). Barcode ligation incubation and stop were done as described above. These steps were repeated 2 more times for a total of 4 barcoding rounds. Finally, the barcoded SPC pool was washed 5 times with SB and was stored at 4 °C with final volume of 200 µl.

### Library preparation

The barcoded SPC pool was divided in 4 PCR tubes with ∼1.250 expected barcoded genomes each (50 µl per tube). Each tube received 2 µl of release reagent (Atrandi) and was incubated at room temperature for 5 minutes to disrupt the SPCs and free the barcoded DNA. The pooled DNA was purified with 0.8X AmPure XP beads (Beckman Coulter) and was eluted in 27 µl of nuclease-free water per tube. DNA fragmentation, adapter ligation and library amplification were done using the NEB Next Ultra II FS DNA Library Prep kit for Illumina (New England Biolabs), with indexing primers UDI5-UDI8 from the Single Microbe DNA Barcoding Kit (Atrandi), and 12 PCR cycles. Final libraries were purified with 0.8X AMPure XP Beads. Of the 4 final libraries, 2 (∼ 2.500 expected cells) were sent for sequencing. Sequencing was done at NovoGene using a NextSeq (Illumina) in high output mode with 150 pair-end reads.

### Cell demultiplexing and read mapping of SPC-scDNA libraries

Demultiplexing of the raw sequencing data into individual single-cell FASTQ files was performed using custom Python scripts in a two-step procedure. Initially, the second read – containing four concatenated 8-nucleotide barcodes with structure [NNNNNNNN]AGAA[NNNNNNNN]ACTC[NNNNNNNN]AAGG[NNNNNNNN]T – was scanned for a perfect match against a list of expected barcode sequences of 8- nucleotide barcodes at the first, second, third, or fourth position. Barcodes were retained only if they perfectly matched the list. Barcodes observed at least 1,000 times were classified as dominant barcodes. The process of screening the second read was then repeated for the identified dominant barcodes, extending the dataset by permitting up to one mismatch per 8nt barcode with the whitelist barcodes. This resulted in paired FASTQ files (R1 and R2) generated for each single cell.

These paired FASTQ files were then aligned to the *Leishmania donovani* BPK282 version 2 reference genome^18^ using BWA^33^. Alignments were filtered for a minimum mapping quality of 30, duplicates were removed using Picard^34^, and only properly paired reads were retained. The resulting BAM files served as input for the bamCoverage module of DeepTools^35^, which calculated average coverage over 20 kb windows normalized to counts per million reads. This was used to generate a count matrix of cells (columns) by bins (rows). Counts were then randomly subsampled to a total of 20,000 reads per cell – matching the number of reads found in the least covered true cell in the scDNA-10X datasets (see below) – to allow cross-sample comparisons. The count matrix was used for cell quality control and karyotyping.

### Single-cell sequencing with 10X-scDNA (10X Genomics)

ScDNAseq data was already available for the BPK081 strain from a previous study using the 10X-scDNA solution^17^. For the HU3 strain, a new experiment was done using a kit bought before the discontinuation of the product. Cell encapsulation, WGA, barcoding and library prep were done as described in the Chromium Single-cell CNV user guide (rev C). Sequencing was carried out at the Genomics Core Leuven on a NovaSeq SP sequencer (Illumina) with 150 pair-end reads. Read demultiplexing, mapping, cell identification and read count matrix generation were done with Cellranger DNA (10X Genomics).

### Compensation of GC bias and mappability

Compensation of GC bias was done by fitting a generalized additive model with bin GC content as explanatory variable and the mean normalized read counts as response variable using the mgcv package in R^36^, after removal of outlier values for GC and count. The resulting model was used to predict expected counts for each bin based on their GC content, and a bin-specific correction factor was computed as the ratio between the median of the total predicted values to the predicted value for each bin. Raw counts were then adjusted by multiplying each bin’s counts across all cells by its respective correction factor. Negative values arising from the correction were set to zero. The corrected count matrix was then used in downstream karyotyping and CNV analysis.

To determine bin mappability, i.e., the fraction of reads that are uniquely aligned to a given genomic bin, a paired-end artificial library of 150 bp reads was simulated with ART Illumina^37^ using a 400 bp insert size, 60 bp standard deviation, and 1x coverage. These reads were mapped back to the BPK282v2 reference genome with BWA-MEM, and the mappability of a bin was calculated as the fraction of reads mapping to it with MAPQ >30 over total mapped reads. Bins with mappability below 0.7 were considered non-mappable and excluded from downstream analyses for karyotyping and CNVs.

### Data quality control

True cells were distinguished from background signal by ranking barcodes in descending order of total read count and plotting the log10-transformed read counts against log10- transformed barcode ranks. The inflection (“knee”) point of this curve, representing the transition from true cells to background, was identified by detecting the point of maximum perpendicular distance from a line connecting the curve’s endpoints. Barcodes to the left of the knee were retained as true cells.

For estimation of coverage evenness, two metrics were used: (i) the gini index, which was calculated for each cell using the gc-corrected and cell-normalized read counts per 20 kb bin with the function ‘gini’ from the ineq package38, and (ii), the median absolute pairwise difference (MAPD) which corresponds to the median absolute difference between the gc-corrected normalized counts across consecutive bins39.

To distinguish replicating (S-phase) cells from non-replicative (G1/G2) cells, we developed an “S-phase score” algorithm based on the observation that cells undergoing genome replication display a wave-like coverage with peak surrounding replication origins, which can cofound karyotyping. For each cell, and for each chromosome, locally estimated scatterplot smoothing (LOESS) model is fitted to the coverage profile (read count per 20 kb bin as a function of genomic bin position). The fitted curve represents the smoothed intra-chromosomal coverage trend. Then, for each bin, we calculated the absolute distance between the LOESS fit and the mean coverage of the corresponding chromosome in that cell. This absolute deviation captures the extent of fluctuation around a flat (non-replicative) coverage profile. We computed the mean absolute deviation per chromosome, and for each cell, selected the five chromosomes with the highest deviations. The final S-phase score for the cell is the mean of these five highest per-chromosome deviation values. High S-phase values indicate structured, wave-like coverage profiles consistent with DNA replication activity, while low S-phase scores reflect flatter profiles characteristic of non-replicating cells. Cells displaying an S-phase score greater than *Q3* + 1.5 x IQR – where *Q3* is the third quartile and *IQR*is the interquartile range – were considered replicative and removed from karyotyping analysis.

Lorenz curves were done for each cell by arranging genomic bins in ascending order for read counts and then calculating the cumulative read count sum. Cumulative sums were normalized by total count in each cell, and Lorenz curves were then summarized across cells of the same sample by calculating the mean cumulative fraction of counts at each genomic fraction.

### Doublet detection

Doublet detection was done as previously reported^17^. In summary, previously generated bulk whole genome sequencing data of strains HU3 and BPK081 was first used to determine strain specific variants. This was done with the Genome Analysis Toolkit (GATK) version 4.1.4.1. This information was then used to attribute reads from the single-cell data to each strain. Cells exhibiting a mixture of diagnostic SNPs from both strains above 5% were classified as inter-strain doublets. Total doublet frequency was estimated under the assumption of random cell pairing, by dividing the observed frequency of inter-strain HU3/BPK081 doublets by the expected proportion of mixed doublets among all doublets, 2*pq*, where *p* and *q* are the relative frequencies of HU3 and BPK081 singlets respectively.

### Single-cell karyotype estimation

Karyotype estimation was done as previously described^17^. In summary, raw somies were determined by averaging the counts of each chromosome and multiplying them by the cell’s scale factor which tries to infer the baseline ploidy of the cell by approximating the normalized values to integers. For this, we iteratively test, for each cell, scale factors within the range of 1.8 to 5 and compute, for each scale factor, the mean absolute distance of the scaled values to the nearest integer. The scale factor that minimized this distance is selected as the optimal estimate of the cell’s ploidy. This method is based on the assumption that true somy values are integers, and that the optimal scale factor should result in values that are as close to integers as possible. Afterwards, raw somy values were converted to integers using Gaussian Mixture Models built at populational level for each chromosome using the mixtools package in R^40^, with one gaussian fitting each possible copy number state for that particular chromosome. The combined integer somy values of a cell defines its karyotype.

### Copy number variations

Copy number variations were determined for two loci which differ between the two strains used in this study: a ∼4 kb locus in chr29 which is present in HU3 but absent in BPK081, and the M-locus, a ∼8 kb locus in chr36 which is present in BPK081 but absent in HU3. To estimate differences in the copy numbers, we generated a new count matrix with DeepTools as mentioned before, but this time with 100bp bins. Correction for GC bias and mappability was done as mentioned above. We then aggregated the corrected counts of the bins inside each locus and divided by the median count of all other bins of the chromosome. This gave an estimated haploid copy number for the locus. Gene-level CNVs were similarly defined by dividing the median read count inside annotated genes by the median count of the chromosome.

### Nucleotide Variants

Additionally, all BAM files were used as input for variant calling with the Genome Analysis Toolkit (GATK) version 4.1.4.1 following the GATK best practices workflow: single-cell level variant calling with HaplotypeCaller generating GVCF files, combining GVCFs of single cells from the same sample using CombineGVCFs, joint genotyping with GenotypeGVCFs, and variant filtering with SelectVariants and VariantFiltration commands according to GATK recommendations. Known drug resistance marker regions were extracted from the VCF files using BCFtools and visualized with the ggheatmap from ggalign^41^ package in R. A variant was considered confidently called when it was covered, at single-cell level, by at least one deduplicated read with Phred quality above Q30 and mapping quality (MAPQ) also above 30.

### Phylogenetic reconstruction

For reconstruction of phylogenetic relationships between cells, the ScisTree2 package was used ^42,43^. First the allele depth matrix generated with GATK was filtered in R with the following criteria: (i) Heterozyogous alleles (frequency < 0.9) were converted to homozygous reference allele, as the package expects only homozygous variants, (ii).

Alternative alleles were only retained if mapped by two or more reads, otherwise they were also converted to reference, (iii), alternative alleles were only maintained if found in 2 or more cells, and (iv) loci with NA values in more than 40% of the cells were removed. The filtered allele depth matrix was then used to estimate genotype probabilities using the probability.from_reads function from ScisTree2 in Python, with default parameters (allelic dropout = 0.2, sequencing error = 0.01, and posterior = True). Trees were constructed with ScisTree2 function, with maximum number of iterations set to 1000. The generated trees were visualized in R using ggtree^44^.

## RESULTS

Two *L. donovani* strains, BPK081, and HU3, were chosen based on the following criteria: (i) HU3 has 122.495 small nucleotide variants (SNVs) compared to the reference BPK282v2 genome, while BPK081 has only 332 variants. This difference is useful to determine doublet rate and the ability to distinguish genotypes with our single-cell approach; (ii) BPK081 is a clonal strain, while HU3 is a not and may therefore have a greater genomic diversity; (iii) both strains differ in bulk aneuploidy profiles, with BPK081 displaying aneuploidy only in chromosome 31, while HU3 presents 12 polysomic chromosomes, and lastly (iv), BPK081 has a known intra-chromosomal CNV in chromosome 36 in a region known as the M-locus which is absent in HU3^25^, while HU3 has a CNV in chromosome 29 which is not detected in the bulk data of BPK081. These differences were used to estimate the method’s ability to distinguish CNVs. Moreover, high throughput scDNAseq data has already been generated for BPK081 in a previous study^17^ using the Chromium single-cell CNV solution from 10X Genomics (referred here as 10X-scDNA), which is considered here as the gold standard technique. Although the datasets came from different cultures derived from the same cryo-preserved sample (Figure 1A), we could compare the single-cell karyotypes determined in the present study against those reported with the 10X-scDNA solution. Here we also performed 10X- scDNA on HU3 for an additional comparison.

**Figure 1.**
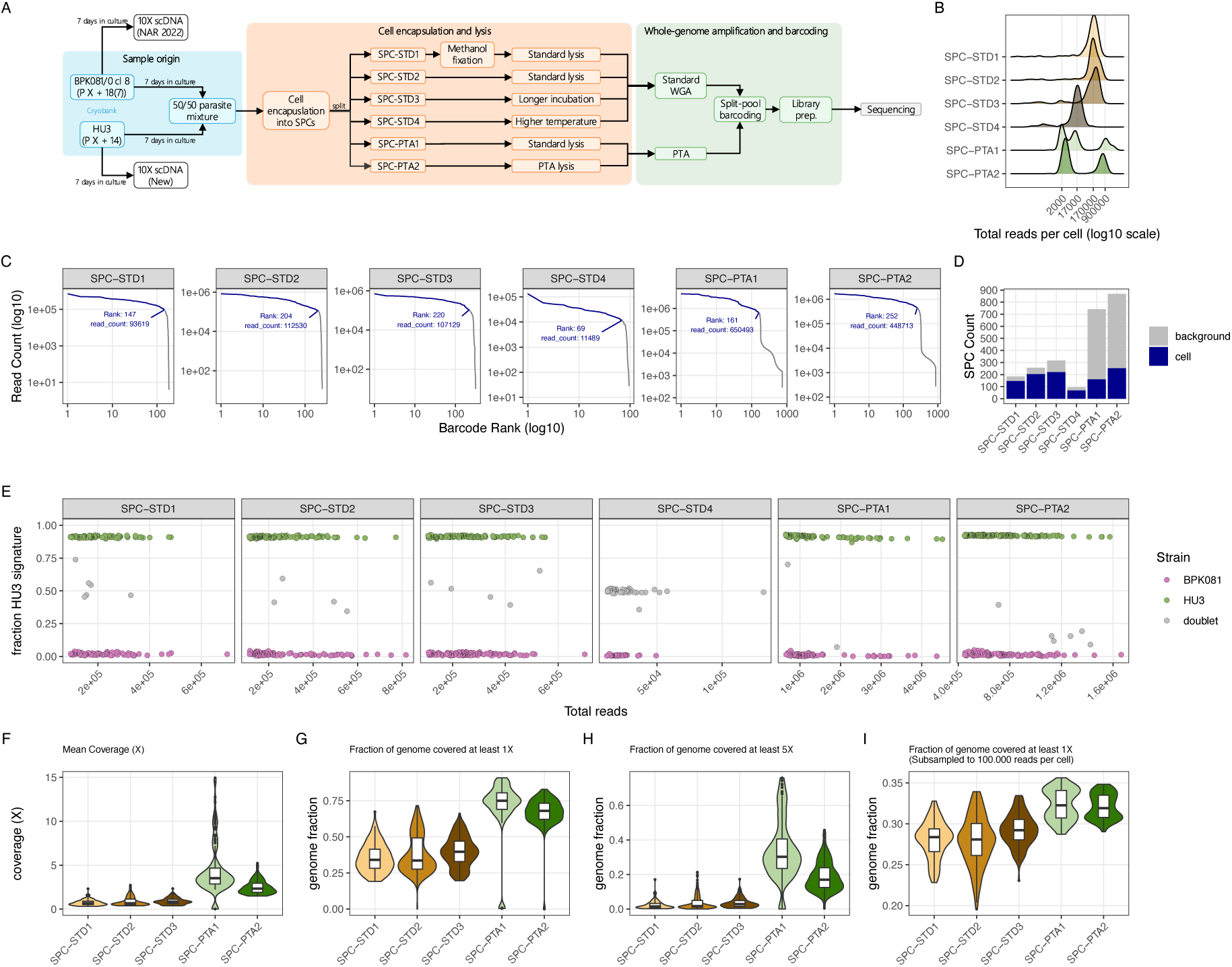
Sequencing metrics and cell characterization in SPC-based scDNAseq. **A:** Schematic representing the experimental setup of the present study. Passage numbers of the two strains are indicated as P X (unknown number of passages) + passages since isolation from clinical sample (18 for BPK081, 14 for HU3), and in the case of BPK081, the number inside parenthesis indicate number of passages since cloning. Standard lysis was done with 0.8M of KOH for 15 minutes at 23 °C. Longer incubation was done for 1h instead of 15 minutes. Higher temperature was done at 9 °C instead of 23 °C. PTA lysis indicates the lysis protocol of the PTA kit (ResolveDNA^TM^ – Bioskryb). **B:** Distribution of the total number of reads associated with each SPC barcode in all 6 samples. X-axis (log10 scale) show regions where some of the distributions peak. **C:** Distinction between ‘true cells’ and background signals. Read counts associated with each barcode were ranked in a log10 space and the ‘elbow’ point in the curve was set as the threshold for background signal. The labels show the rank and read count of the last SPC considered as ‘true cell’ **D:** Number of SPCs containing a ‘true cell’ (blue) or a background signal (grey) in each sample. **E:** Evaluation of HU3/BPK081 doublets (grey dots) in each sample. Y-axis show the percentage of variant reads with HU3 signature, which was predetermined based on bulk whole genome sequencing of HU3. **F-I**: analysis of nucleotide coverage depth in each sample. Coverage (F) is defined as the average number of reads mapping each nucleotide in the genome. Figures G and H show the percentage of the genome that was covered by at least 1 and 5 reads respectively, while figure I show the percentage of the genome covered by at least 1 read after cells were normalized to 100,000 reads per cell. Violin plots display the distribution of the data (kernel density), with embedded boxplots indicating the median (thick line), the interquartile range (IQR – shown by a box spanning the 25th to 75th percentile), and whiskers (lines) extending to 1.5×IQR. Points beyond the whiskers indicate outliers.

### scDNAseq with semi-permeable capsules

We based our scDNAseq approach on the Single Microbe DNA Barcoding Kit from Atrandi Biosciences^TM^, which uses semi-permeable capsules (SPCs) for high throughput cell encapsulation^30^. The kit includes a pre-defined cell lysis protocol developed for bacteria, which starts with a methanol fixation step followed by incubation for 15 minutes at room temperature in the provided lysis buffer (0.8 M KOH, 20 mM EDTA, 200 mM DTT). Because it was unclear whether this protocol would be effective for *Leishmania* cells, we also tested modified lysis conditions by either extending the incubation time from 15 to 60 minutes or increasing the incubation temperature to 99 °C. In addition, we assessed if the protocol could be simplified by omitting the methanol fixation step. These modifications were first evaluated in pools of ∼1,000,000 parasites per condition, by measuring with qPCR the quantity of free-floating DNA in the supernatant after parasite lysis and centrifugation. This analysis indicated that omiting methanol fixation in fact improved DNA release, whereas no clear differences were observed among the different lysis conditions (see Supplementary Text, Supplementary Figures 1 and 2, and Supplementary Tables 1 and 2).

In the next step, we tested these different lysis conditions in the full scDNAseq protocol. For that, a 50/50 BPK081/HU3 mixture was submitted to SPC encapsulation, with trapped cells being visualized with SYTO5 staining. As expected, most SPCs appeared empty, with the remaining ones containing only a single parasite (supplementary figure 3A). No SPC containing multiplets were found by microscopy inspection. This also confirmed that cells remained alive after encapsulation (supplementary video 1). The generated SPC pool was then divided in 6 samples to test different lysis and WGA conditions. Because the Single Microbe DNA Barcoding kit uses a WGA method based on MDA, a method which failed to recapitulate structural variants in a previous work^29^, we decided to test PTA as an alternative WGA due to its reported improved coverage evenness ^31^. A summary of the different conditions and general statistics of each sample is provided in Table 1.

After lysis and WGA, SPCs were stained again with SYTO5 to detect amplified DNA. Strong SYTO5 staining were seen in all samples except for sample SPC-STD4, which had no detectable signal, indicating WGA failure (supplementary figure 3B). SPC-STD4 also exhibited enlarged SPCs, which could indicate they were damaged during lysis. Moreover, in sample SPC-STD1, although positive SPCs were entirely populated by amplified DNA, fixed parasites were still visible on some of them (supplementary Figure 3C), indicating that the methanol fixation led to partial lysis.

After confirmation of successful WGA, SPCs were pooled, barcoded and sequenced. Sequencing output was first assessed by read counts per sample (Figure 1B). Standard WGA samples showed a unimodal distribution around ∼170,000 reads per cell, except sample SPC-STD4, which peaked at ∼17,000 reads, consistent with its observed WGA failure. In contrast, PTA samples displayed bimodal/multimodal distributions, with a major peak at 600,000–900,000 reads, suggesting stronger cell DNA amplification, and minor peaks between 2,000–17,000, likely from amplification of background DNA.

To separate true cells from background signal, we applied a barcode rank approach^45^. All samples exhibited a characteristic sharp drop-off in read counts, indicating a clear inflection point that distinguishes cell-associated barcodes from background (Figure 1C). The PTA samples contained the cells with highest sequencing depth, with thresholds at 656,209 reads (SPC-PTA1) and 448,715 reads (SPC-PTA2). Thresholds for the standard WGA samples were lower (109,851 - 180,703 reads in samples SPC-STD1- 3) and lowest in sample SPC-STD4 (11,413 reads), which also had the fewest SPCs flagged as true cells (70 SPCs). PTA samples had more total detectable SPCs, but a similar number of true cells (160 and 253 in SPC-PTA1 and SPC-PTA2) compared to standard WGA (135, 206, and 166 in SPC-PTA1-3 respectively) (Figure 1D and Table 1). Only true cells were retained for downstream analysis. We then assessed the presence of doublets by quantifying HU3-specific variants in each SPC, using a curated set of HU3- specific SNPs identified from bulk genomic sequencing. Successful WGA samples showed only 2–6 inter-strain doublets (4-12 estimated total doublets), while most SPCs in sample SPC-STD4 contained a mixture of BPK081 and HU3 reads (Figure 1E and Table 1), suggesting that amplicons originated from mixed free-floating DNA. According to follow-up guidance from the kit technical support team, it was suggested to use a shorter incubation time (10 min rather than 15 min) and a lower KOH concentration (0.2 M rather than 0.8 M) for heat-based lysis. We therefore infer that the combination of prolonged incubation, high temperature, and high KOH concentration in SPC-STD4 damaged the SPCs and promoted DNA leakage before WGA. Because of the clear failure in WGA and the lack of single-cell signal, sample #SPC-STD4 was excluded from downstream analysis.

Nucleotide coverage was also higher in PTA samples (Figure 1F), with ∼67-71% of the genome being covered by ≥1 read (Figure 1G), and ∼18-33% by ≥5 reads (Figure 1H), while samples SPC-STD1-3 showed 35–39% coverage at ≥1 read and 2.4–3.5% at ≥5 reads. After equalizing each sample to 100,000 reads per cell (the number of reads observed in the least covered true cell in sample SPC-STD1), PTA still performed best, with ∼32% of the genome being mapped at least once, while in the standard WGA this value was ∼27– 29% (Figure 1I). In addition, no conclusive difference was seen across the different lysis conditions. Together, these results indicate that i) methanol fixation can be omitted for parasite lysis, ii) the standard lysis conditions of both the single microbe DNA barcoding and the ResolveDNA kits work well with *Leishmania*, and iii) PTA achieves higher yields of DNA amplification than standard WGA.

### Evaluation of coverage evenness

Detecting structural variants such as CNVs and aneuploidy is a key application of scDNAseq and requires uniform genome coverage to ensure that read counts reflect true copy number changes. Thus, we compared coverage uniformity in our dataset with that of two libraries generated using the Chromium Single-Cell CNV solution (10X-scDNA), considered the gold standard for this application. For this, we first subsampled each cell to 20,000 reads, matching the cell with the least read count in the 10X-scDNA dataset. After correcting for GC bias (supplementary figure 4), a Lorenz curve was made to evaluate global read distribution, indicating that the PTA samples had a similar profile compared to the 10X-scDNA libraries, while samples SPC-STD1-3 showed a skewer distribution (Figure 2A). This also reflected in lower gini coefficients compared to SPC- STD1-3 samples as well as lower MAPD scores, achieving values comparable to the 10XscDNA libraries (Figure 2B-C). We also introduced the S-phase score, which is used to identify local intra-chromosomal biases which might skew the somy estimation, common in cells under S-phase but also affected by coverage evenness. The PTA samples displayed similar S-phase scores compared to the 10X-scDNA libraries, while samples SPC-STD1-3 had consistently the highest (Figure 2D). Importantly, no clear distinction was observed regarding lysis conditions. The improved uniformity in PTA was also noticeable when visualizing the normalized read coverage across all genomic bins (Figure 2E-G). In the standard WGA samples, coverage profiles were considerably noisier, with uneven coverage across cells that obscured clear aneuploidy patterns (Figure 2F). In contrast, the PTA samples (Figure 2G) exhibited much smoother profiles, with well-defined aneuploidy regions that closely resembled the high-quality profiles obtained with the 10X-scDNAseq reference (Figure 2E). Together, these analyses demonstrate that PTA improved coverage uniformity, achieving performance comparable to the 10X-scDNAseq gold standard.

**Figure 2.**
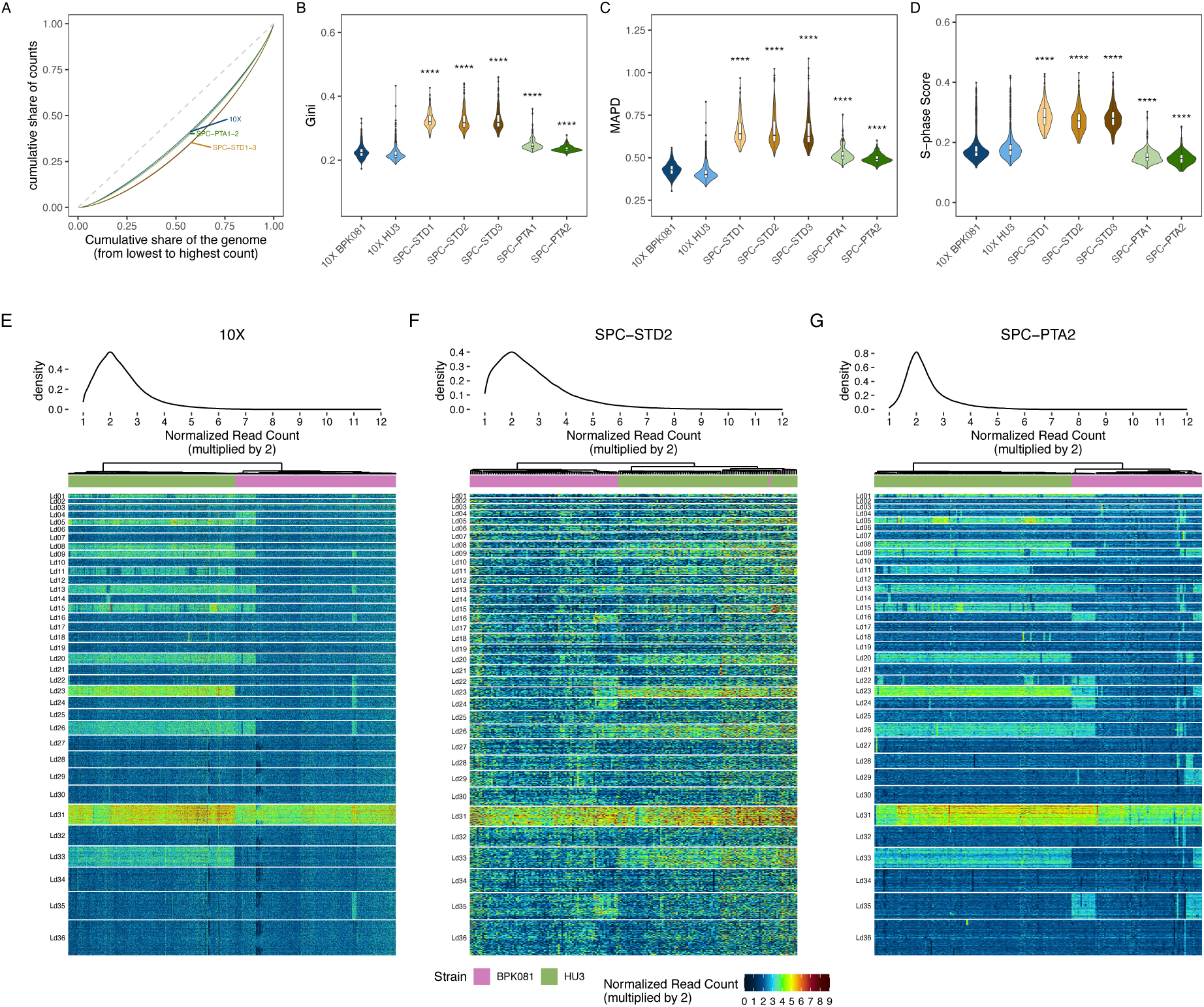
Evaluation of the impact of the different WGA methods on coverage evenness. **A:** Lorenz curves showing the cumulative fraction of sequencing counts as a function of the cumulative fraction of the genome covered in the scDNAseq datasets. The dashed diagonal represents perfect uniform coverage. Curves deviating downward from the diagonal indicate increasing coverage unevenness across the genome. The blue line indicates the 10X-scDNA datasets, while the green lines show samples SPC-PTA1-2, and orange lines represent samples SPC-STD1-3. **B-D:** violin plots showing the gini (B), MAPD (C), and S-phase (D) scores for each dataset compared to the 10X-BPK081 and 10-HU3 libraries. Asterisks indicate the p-values of a Wilcox test of all samples against both 10X-scDNA libraries combined adjusted with the Hochberg method. **** = p-value < 0.0001. **E-G**: heat map showing the read count of each 20k bin (y axis) in each cell (x – axis) in each chromosome (row groups). Colors indicate the read count of each bin normalized by the median read count in the cell multiplied by 2. Cells were arranged with the ward.D2 hyerarchical clustering algorithm based on the heat map values. Annotation bars indicate the corresponding parasite strain. A density plot on top shows the distribution of values in the heat map. Here sample SPC-STD2 is shown as a representative of standard WGA samples, while sample SPC-PTA2 represents the PTA samples.

### Single-cell Karyotyping

To define the impact of the improved coverage uniformity on chromosome copy number estimation, we first checked the distribution of raw somy values, i.e., the normalized average read count of chromosomes multiplied by the cells’ scale factor. In the SPC-STD datasets, we observe a more continuous distribution, with a substantial number of values falling between integers (Figure 3A), likely due to the greater noise seen in this data. Conversely, PTA produces a markedly discrete distribution, with sharp peaks centered on integer values (e.g., 2, 3, and 4), and relatively few intermediate values (Figure 3B). Since chromosome copy numbers are inherently integers, PTA’s output aligns more closely with biological expectations. This also reflected into less cells being scaled to ploidies higher than 2, as noisy coverage seen in standard WGA distorted depth ratios and consequently led to incorrect scale factors. Cells scaled to ploidies higher than 2 in PTA could represent true polyploid cells which can spontaneously arise in culture^46^, as well as cells at S/G2 phase.

**Figure 3.**
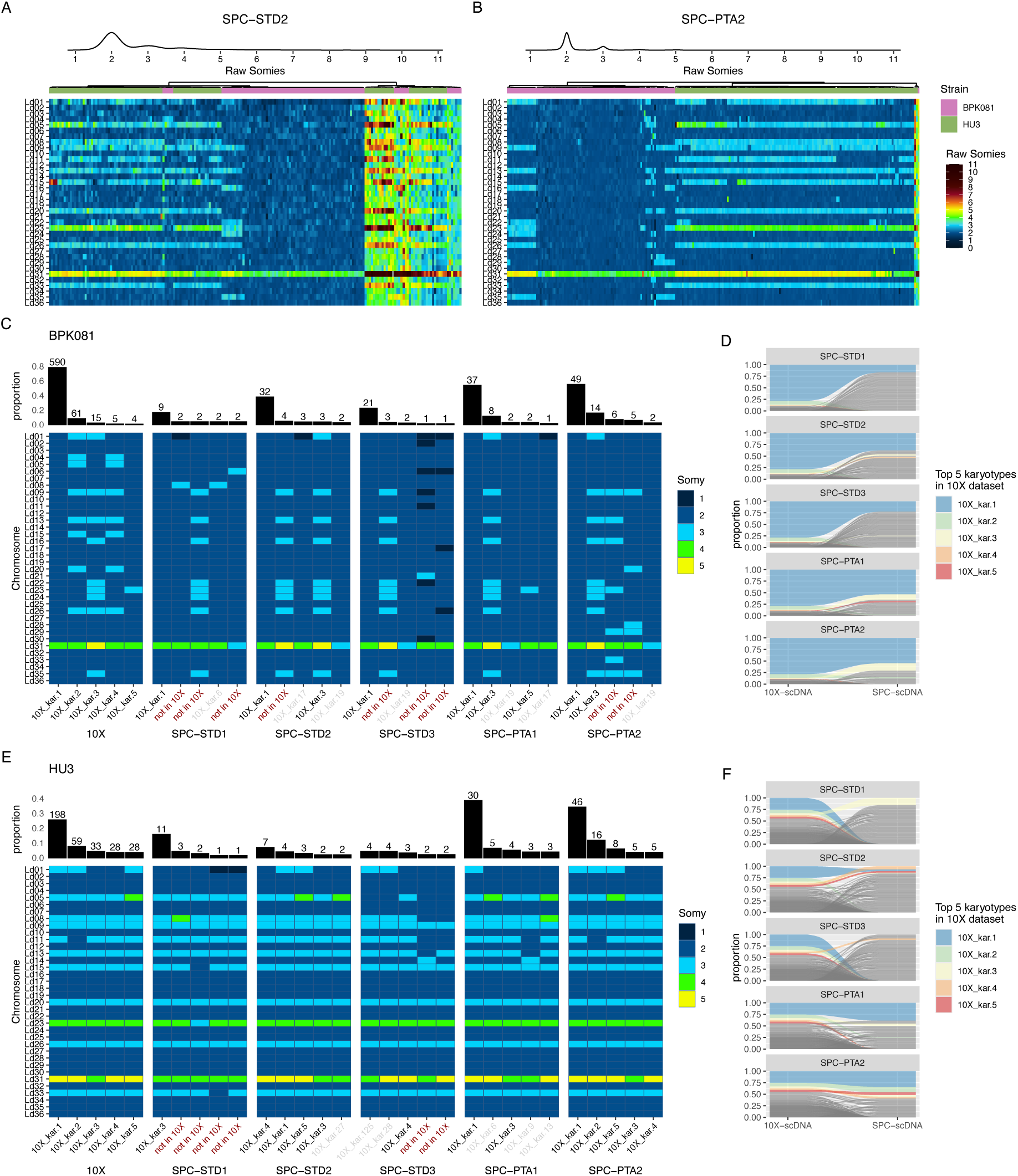
Evaluation of the standard WGA and PTA for determination of chromosome copy numbers in single-cells. **A- B:** Heat maps showing the estimated raw somy values for each cell in sample SPC-STD2 (A) SPC-PTA2 (B). Raw somy values were calculated as the mean count of each chromosome (normalized by the cells mean) multiplied by the cell’s scale factor. Cells were arranged with the ward.D2 hyerarchical clustering algorithm based on the heat map values. A density plot on top shows the distribution of values in the heat map. Annotation bars indicate the corresponding parasite strain. **C:** heat maps comparing the 5 most frequent karyotypes identified the BPK081 cells of each in the SPC- scDNA samples against the 10X-scDNA reference. Heat map colors indicate the integer copy number of each chromosome (y-axis) in each of the 5 karyotypes (x-axis). Columns on top indicate the proportion of the total sequenced population displaying each karyotype. Numbers on top of the bars show the number of cells instead. **D:** Sankey diagram showing the proportion (y-axis) of each BPK081 karyotype in the SPC-scDNA samples compared to those found in the 10X-scDNA dataset. Each karyotype is represented by a line which width indicates its proportion and height indicating its position. The top 5 karyotypes found in the 10X-scDNA reference are colored. **E-F**: The same as C-D but for the HU3 cells.

Then, we compared the karyotype profiles of the SPC datasets with the 10XscDNA libraries of each respective strain. The karyotype profiles obtained from PTA samples were largely consistent with those generated by 10X-scDNA, whereas standard WGA samples showed poorer reproducibility. For the BPK081 strain, the dominant profile where only chromosome 31 was aneuploid was the most frequent in all samples, representing ∼60% of cells in PTA samples, in close agreement with the ∼80% reported in the 10X-scDNA dataset (Figure 3C). The highly aneuploid karyotypes 2 and 3 seen in 10X-scDNA were also detected in PTA samples 5 and 6, but not in samples SPC-STD1–3, likely due to underestimation of the chromosome 1 trisomy (Figure 3C-D). Importantly, the SPC-scDNA and 10X-scDNA datasets originated from different cultures derived from the same cryopreserved population, so some variation in karyotype content was expected, especially given the instability of the BPK081 dominant karyotype in culture^21^. Accordingly, an even more similar pattern was observed for the HU3 strain: PTA reproduced the same dominant karyotypes identified with 10X-scDNA with a remarkable similarity in proportions between them, while standard WGA produced divergent and inconsistent profiles (Figure 3E–F). Altogether, these results demonstrate that PTA- based amplification preserves karyotypic structure with high fidelity, enabling more accurate reconstruction of chromosome copy number profiles than the standard WGA.

### Detection of small structural variants

To test whether we could detect smaller structural variants, we took advantage of the existence of two different CNVs which are unique to each strain – a ∼8 kb CNV (M-locus) in chromosome 36 unique to BPK081, and a ∼4 kb CNV in chromosome 29 characteristic of the HU3 strain. For this analysis, we increased the resolution to 100 bp bins, using the chromosome-normalized median read count of the bins inside the CNV as an estimation of haploid copy numbers. In the PTA samples, all BPK081 cells displayed a clear and consistent increase in coverage depth at the M-locus compared to HU3, in contrast to the standard WGA samples where only a fraction of BPK081 cells had a detectable amplification of the M-locus (Figure 4A). Likewise, the CNV in chromosome 29 was consistently detected in all HU3 cells from PTA samples, with no BPK081 cell displaying them, while with standard WGA only a fraction of HU3 cells had a detectable amplification (Figure 4B). The distribution of estimated haploid copy numbers of both CNVs shows sharp, well-defined peaks at expected integer copy numbers for the SPC- PTA samples, with all BPK081 cells showing amplification at the M-locus and all HU3 cells showing amplification at chromosome 29 (Figure 4C). By contrast, the standard WGA samples (SPC-STD1–3) exhibited broad and noisy distributions, with many intermediate values and amplifications detected only in a subset of cells. Interestingly, in PTA samples, minor peaks at alternative integer values were observed for both loci, which could indicate the presence of subpopulations with distinct CNV states.

**Figure 4.**
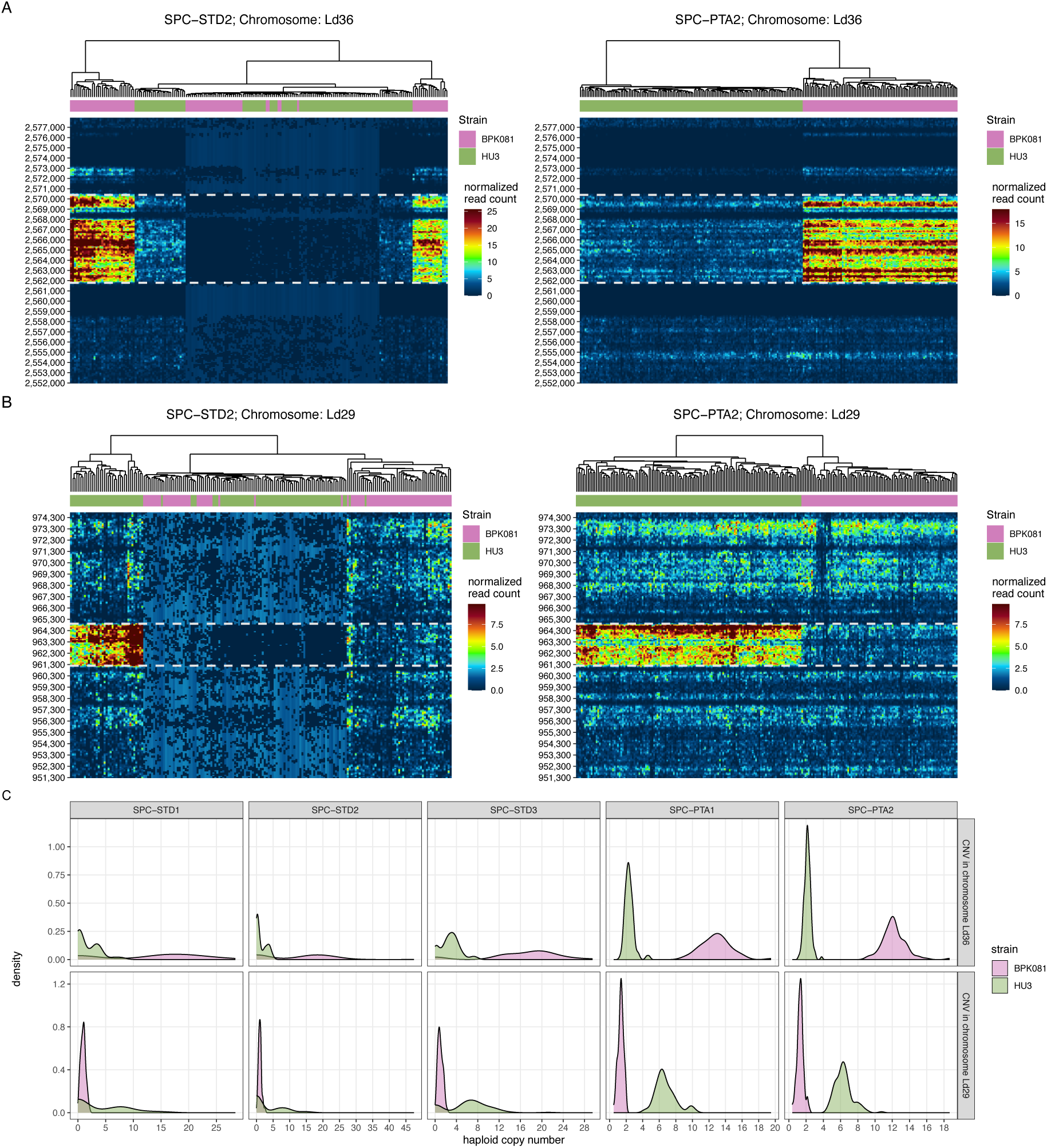
Detection of strain-specific CNVs. **A:** heat maps showing the read counts of samples SPC-STD2 (left) and SPC-PTA2 (right) across 100bp bins in the region surrounding the m-locus in chromosome 36. Heat map colors indicate normalized read count for each bin. Cells were arranged with the ward.D2 hierarchical clustering algorithm based on the heat map values. Annotation bars indicate the parasite strain. The white dashed lines highlight the known CNV region. **B:** same as A, but showing the normalized read counts for the CNV found in the chromosome 29. **C:** Distribution of the estimated haploid copy number of the m-locus (top row) and the CNV in chromosome 29 (bottom row) in each cell and in each sample. Haploid copy numbers were estimated by dividing the average count in the bins located in the m-locus by the average count of the other bins of the chromosome. Colors indicate the strain.

We also investigated if other CNVs could be present in our data using a gene-level unsupervised approach. For that, we divided the median read count of each annotated gene by the median count of the chromosome and looked for genes with median value 1.5 times higher than the chromosome median (supplementary figure 5). As expected, The SPC-PTA samples had substantially higher sequencing depth, making genome-wide CNV calculation feasible. Notably, the two previously identified loci - the M-locus on chromosome 36 in BPK081 and a known CNV on chromosome 29 in HU3 - stood out clearly against this background, consistent with the CNVs we had already characterized by bulk genome sequencing. However, this showed no clear, consistent signal across genes except for these two previously identified loci. Still, these results confirm that PTA retains better small structural information.

### Nucleotide sequence variants

To investigate the applicability of our method in detecting heterogeneity at nucleotide sequence level, we first assessed its ability to distinguish the two *L. donovani* strains in a *de novo* way, i.e., without prior bulk sequencing data. This would mimic the scenario of samples with mixed strain infections. Principal component analysis (PCA) based on nucleotide variants showed a clear distinction between cells from HU3 and BPK081 strains in all samples (Figure 5A), indicating that both the standard WGA and PTA methods are good enough to separate parasite populations of the same species.

**Figure 5.**
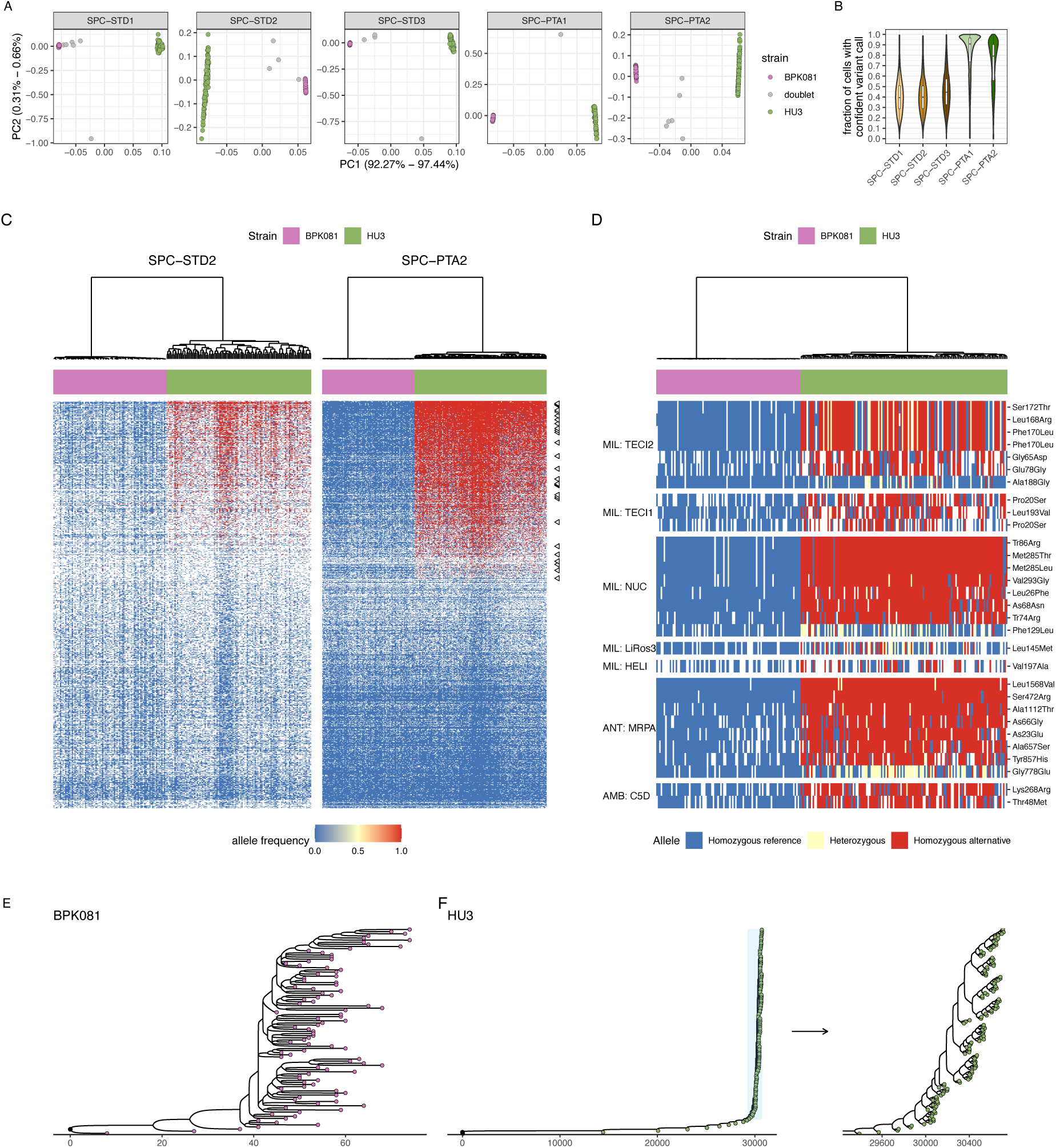
Identification of nucleotide variants with the SPC-scDNA workflow. **A:** Principal component analysis (PCA) based on nucleotide variants identified in each single-cell dataset. PCA was done using PLINK2 after removal of linked variants. Colors of the dots indicate the strain. **B:** Violin plots indicating the fraction of cells in which each variant was confidently called (i.e., covered, at single-cell level, by at least one deduplicated read with Phred quality above Q30 and mapping quality (MAPQ) also above 30). Included boxplots indicate the median (thick line), the interquartile range (IQR – shown by a box spanning the 25th to 75th percentile), and whiskers (lines) extending to 1.5*IQR of the data. **C:** heat map showing the estimated allele frequency of 1000 randomly sampled variant loci in samples SPC-STD2 (left) and SPC-PTA2 (right). Color scale goes from blue (allele frequency of 0) to yellow (0.5) and red (1). White indicates missing values (NAs). Annotation bars indicate the cell strain, and cells were arranged with the ward.D2 hierarchical clustering algorithm based on the heat map values. The black arrows to the right side of the heat map of SPC-PTA2 show the drug resistance-associated loci. **D:** A zoom in the drug resistance-associated genes. Here only genes showing heterogeneity in nucleotide sequence are shown. The TEC1, TEC2, NUC, LiRos3, and HELI genes are associated with resistance against miltefosine (MIL), while MRPA is associated with Antimony (ANT) resistance, and C5D is related to amphotericin B (AMB). The right row labels indicate which type of mutation each variant is predicted to cause in the protein expressed by those genes. **E-F**: Phylogenetic trees built based on homozygous variants found in the dataset of SPC-PTA2. Trees were constructed with the ScisTree2 pipeline for the BPK081 strain (E) and HU3 (F). In F, an inset displays an expansion of the extreme right part of the tree. Numbers in the x-axis indicate the number of different variants between nodes.

Next, we investigated if we could detect low-frequency nucleotide polymorphisms at particular loci of interest with single-cell resolution. We performed a global search for variants in all datasets and observed a greater proportion of confidently called variants (variants covered by at least 1 read at single-cell level after read filtering) in the PTA samples, where variants were resolved in ∼80-90% of the cells on average, compared to standard WGA (∼60 %) (Figure 5B). This is also noticeable when visualizing the estimated allele frequencies for each variant loci (Figure 5C). Because of the high prevalence of missing values in the standard WGA samples, we moved forward only with the PTA samples.

As proof-of-concept, we examined genes known to contribute to reduced susceptibility to leishmanicidal drugs to investigate potential presence of heteroresistance in the parasite population. We found a relative homogeneous genotype in BPK081, with few cells displaying heterozygous single nucleotide polymorphisms (SNPs) in some of the analyzed genes (Figure 5D), which is in accordance with the fact that this strain is a clone with relative few passages (7 since cloning). Conversely, HU3 cells displayed more diversity in these same loci, with small subset of cells displaying homozygous SNPs in genes related to resistance to antimony, miltefosine, and amphotericin B. This highlights the presence of spontaneous variants with potential clinical relevance which would likely have been missed by bulk sequencing.

Lastly, we used the nucleotide information to infer the short-term evolutionary relationship between cells along propagation in culture. The resulting tree revealed a compact structure with limited branch lengths and a maximum divergence of fewer than 70 nucleotide variants between cells in the BPK081 (Figure 5E). This restricted diversity indicates that the clonal strain retained a largely homogeneous genetic composition, with only minor sequence changes detectable across individual cells, in accordance with the fact that BPK081 is a clone with limited expansion time in culture. In contrast, the HU3 strain, which was never clonally derived and has been maintained for at least 15 passages (but likely many more passages), displayed higher heterogeneity (Figure 5F). The tree spanned over 30,000 nucleotide variants, with a small subset of 8 cells forming long branches, followed by multiple subclades emerging within the population. A magnified view of the terminal region of the tree further highlights the presence of distinct subgroups differing by a few hundreds of variants, likely arising during ongoing diversification in culture.

To determine whether the presence of 8 cells in long branches observed in HU3 in sample SPC-PTA2 could be driven by technical artefacts (e.g., reduced coverage, less variants called), we closely inspected the coverage values in these cells compared to the other cells in the same dataset. We did observe a consistent tendency toward lower coverage and higher variant missingness (undefined loci) in these 8 cells, but still within the range of variation seen across the other HU3 cells (supplementary figure 6A). Importantly, other cells show comparable or even higher levels of missingness/reduced coverage but remain clustered with the main bulk rather than forming long branches in the phylogram. Thus, it is unlikely that missingness/coverage is the sole cause of their phylogenetic separation. This is further supported by a similar analysis in sample SPC-PTA1, which represents and independent sampling of the same parasite population. Indeed, the phylogenetic tree of this sample showed the same overall pattern, where only a small subset of cells was present in long-branches (supplementary figure 6B). However, in the case of SPC-PTA1 these cells did not show lower coverage or higher missingness compared to other cells (supplementary figure 6C). Thus, these results are most consistent with minor divergent subpopulation(s) persisting during long-term culture. As HU3 is a non-cloned isolate, polyclonality remains a plausible explanation for a small subset of more divergent cells. At the same time, we cannot exclude that locus-/depth-dependent calling sensitivity may contribute to the magnitude of inferred mutations in this subset.

## DISCUSSION

In the present study, we developed and validated a high-throughput single-cell genomic approach using *Leishmania* as a model eukaryote. Using SPCs for high throughput cell encapsulation and PTA for WGA, we could simultaneously resolve karyotypes, detect intra-chromosomal CNVs, uncover low-frequency genotypes, and explore population structure. These findings represent a significant advance in dissecting genome instability and adaptation in eukaryotic pathogens at single-cell resolution.

### Improved Whole-Genome Amplification with PTA vs Standard WGA

*Leishmania* display a highly segmented genome: its small size of 33 Mb is divided among 36 chromosomes^47^. This characteristic makes scDNAseq particularly challenging given that WGA methods are usually validated in mammal cells, with nuclear genomes about 100 times larger than *Leishmania*. Thus, to obtain enough material for sequencing, *Leishmania* DNA must be amplified orders of magnitude more compared to a mammalian counterpart, increasing the risk of artefactual biases. Our single-cell genome analysis of *Leishmania* applied primary template-directed amplification (PTA) to overcome the biases associated with other WGA methods such as MDA. PTA was developed as a modification of MDA by introducing exonuclease-resistant terminator nucleotides which limit amplicon length and prevent reamplification of daughter molecules, changing the amplification kinetics from exponential to quasi-linear^31^. This dramatically reduces allelic dropout and amplification bias, thereby preserving the true allelic and structural representation of the single-cell genome^31^. Notably, this method has never been tested in microbial eukaryotes such as *Leishmania*. Here, we demonstrated that *Leishmania* genomes amplified with PTA yielded consistent coverage across chromosomes and between replicates, outperforming the standard WGA – which is based on MDA – in all metrics. When benchmarking against an established 10X Genomics single-cell CNV dataset, the PTA-based workflow reached comparable coverage evenness and recapitulated expected karyotypes more in accordance with the expected distributions, whereas the standard WGA introduced major artifactual karyotypes not seen in the reference data. Altogether, our results demonstrate that PTA’s uniform amplification preserves genomic structure far more faithfully, enabling more accurate reconstruction of chromosome copy number and sequence variants even from single cells with a small genome.

### SPCs as a viable tool for high throughput implementation of PTA

Although PTA offers good performance in WGA fidelity, its application has been limited to low throughput workflows, using cells isolated in plates which are manually processed for WGA in individual reactions ^31,48,49^. Here we applied SPCs as an alternative solution for cell encapsulation to increase throughput. SPCs offer a more flexible alternative to more commonly used single-cell compartimentalization techniques like emulsion droplets, due to its semi-permeable nature, which allows the easy adaptation of multi-step protocols to work with single cells. This allowed us, for instance, to easily change the cell lysis conditions and WGA reactions in our experiment. It also allows the parallel isolation and sequencing of up to 10.000 cells using relatively low-cost instruments^50^, increasing throughput and thus reducing costs. Moreover, the split and pool barcoding strategy can also be exploited as a sample multiplexing system as was done here, by allocating cells of different samples in specific pre-defined wells in the first barcoding round, reducing costs even further when dealing with multiple samples.

With SPCs we could apply PTA to a total of 399 cells (samples SPC-PTA1-2) while using the equivalent of 17 PTA reactions (17 cells if isolated in plates). Considering the costs of the PTA (∼ €3100 for 96 WGA reactions) and library prep kits (∼ €3100 for 96 libraries), a full plate-based system for 96 cells would have costed ∼ €65 per cell excluding sequencing cost. In our hands, this cost was reduced to ∼ €3,9 per cell including the required SPC generation kit (∼ €230 for 1 encapsulation run) the PTA reactions (∼ €550 eur for 17 reactions) the barcoding and library prep kits (∼ €660 for 2 samples in the barcoding plate if using 6 sample multiplexing) and library prep (∼ €130 for 4 final libraries). We also estimate that using the maximum capacity of 10.000 cells would further reduce price per cell to ∼€0,39, although this comes with the caveats of a higher doublet rate and higher sequencing costs to achieve robust per-cell coverage. Overall, SPCs improves scalability and cost-efficiency, making high-throughput single-cell genome sequencing more accessible.

### High-Resolution Karyotyping and CNV Detection

With the combination of SPCs and PTA, we achieved high-resolution single-cell karyotyping of *Leishmania* that surpasses prior methods. Each parasite’s complete karyotype could be determined, and we could even resolve CNVs as subtle as a few kilobases. For example, in a mixed population of two *Leishmania* strains, our method could distinguish two known CNVs – spanning 4 kb to 8 kb in size – which were unique to each strain. Detecting such fine-scale CNVs at the single-cell level is a notable advance.

Traditional single-cell CNV platforms like 10x Chromium yield copy number profiles at sparser resolution – on the order of ∼2 Mb for mammalian cells, and at least ∼200 kb for *Leishmania*^17^ – and typically require pooling signals across multiple cells to detect smaller events^51,52^. In contrast, the high depth and uniformity of our PTA-amplified genomes enable detection of gene-level amplifications or deletions within individual cells. This could be particularly relevant in the context of genomic surveillance of drug resistance markers such as the amplification of the H-locus and M-locus, two CNVs found in some *L. donovani* strains which were linked to antimony resistance in the Indian Subcontinent^19^. Moreover, CNVs have been linked to drug resistance in other pathogenic protists, such as *Plasmodium falciparum*, where CNVs affecting the plasmepsin genes are linked to resistance to piperaquine resistance^53^, highlighting the potential relevance of the method in other organisms.

### Heterogeneity of nucleotide sequence

A further complication in *Leishmania* research is the possible occurrence of mixed infections and within-host diversity. Field studies have shown that individual hosts (human or animal) can harbor multiple *Leishmania* genotypes or even different species simultaneously, with a potential impact on clinical outcome ^26–28^. Moreover, population-genomic studies of the *L. donovani* complex have identified isolates with allele-frequency patterns consistent with mixtures of genetically similar subclones, including mixtures within the same phylogenetic group^54^. Mixed-genotype infections are likely under-detected by routine diagnostics^55^, and confounds bulk genomic analyses. This was evident when cultured parasites showed genomic signatures distinct from their clinical counterparts, likely due to the expansion of initially undetectable genotypes in culture^56^. Similar challenges have been described in other protozoan parasites such as *Plasmodium falciparum*, where polyclonal infections can contain mixed drug-sensitive and drug-resistant genotypes^57,58^ and conventional bulk genotyping may reduce this complexity to binary calls^59^, motivating the development of approaches that resolve subpopulation structure more explicitly. This complexity makes it challenging to link genotype to phenotype and to study evolutionary dynamics, especially for traits like drug resistance or virulence where minor variants can be clinically relevant.

Here we demonstrated the feasibility of distinguishing genotypically different cells by using artificial mixtures between two *L. donovani* strains. Cells were clearly separated without any prior knowledge of the genotypes composing the cell mixture. Moreover, although the HU3/BPK081 mixture represents a relatively divergent two-strain model, the result shown in Figure 5D, which uses only a subset of 30 SNPs, indicates that separation can be achieved even with a limited set of informative loci, and the within-strain structure observed in the BPK081 tree in Figure 5E suggests sensitivity to relatively small mutational differences. This is of particular interest in cases where mixed infections happen between closely related parasite populations^54^. While sequencing single cells directly from host tissues may currently not be feasible, this method could be applied to fresh isolates (after only one or two passages) to detect circulating low-frequency genotypes, distinguish mixed infections from hybrids, and to monitor resistance-associated variants.

As a proof-of-concept, we searched our data specifically for nucleotide polymorphisms in known drug resistance-associated genes. We revealed high heterogeneity in SNPs in a few subsets of genes in the HU3 strain, which constitutes so far undetected/hidden levels of genomic heterogeneity, potentially already present in the parasite infecting the human host. Such within-population variability in drug resistance markers might represent hallmarks of heteroresistance, where genetically distinct subpopulations coexist with differing levels of drug susceptibility^60,61^. Indeed, experimental selection of miltefosine resistance revealed that parasite adaptation to the drug was driven mainly by pre-existing mutations in a miltefosine transporter gene which were undetected in bulk genome sequencing^20^. In addition, another advantage of the single-cell approach is that it enables detecting co-existing mutations in the same cell, as illustrated in Figure 5D. This is particularly relevant in protozoan parasites, where resistance may depend on multi-locus haplotypes or combinations of mutations, as illustrated by resistance against sulfadoxine-pyrimethamine treatment in *Plasmodium falciparum*, which is characterized by mutations in dhfr and/or dhps genes, including triple and septuple mutant forms as well as mutations in both genes^62–64^.

Lastly, we also used the nucleotide information to infer the evolutionary relationship between cells of the two strains, showing that the clonal strain BPK081, maintained for only seven passages, exhibited minimal divergence among individual cells, while a higher diversity was observed in HU3. The ability to reconstruct phylogenetic trees at the single-cell level provides a powerful tool to study cellular evolution in both experimental and natural populations. For instance, using genetically introduced cellular barcoding, our group could track the clonal adaptation to two leishmanicidal drugs *in vitro*, revealing polyclonal origins for the genetic alterations associated with adaptation^20^. Similar information could be gathered with scDNAseq without the need for genetic engineering. More broadly, single-cell phylogenies have proven instrumental in understanding tumor evolution and drug resistance in cancer^65,66^. By resolving genetic lineages at cellular resolution, our method opens opportunities to monitor how eukaryotic cells diversify under selective pressures, weather in protozoan pathogens, or potentially in model systems or human disease.

### Future perspectives

The SPC-PTA scDNAseq workflow described here opens new possibilities for understanding the biology and adaptation mechanisms in *Leishmania* and other microbial eukaryotes. For instance, scDNAseq has been used to study DNA replication programs, revealing fork progression speeds and replication timing along the cell cycle in mammalian cells^67,68^. Unlike most eukaryotes, *Leishmania* has a unique DNA replication program based on a single large replication origin per chromosome which is highly active during S-phase and which is complemented by thousands of minor low- efficiency origin sites that are active during G1 and G2/M phases^69,70^. However, it is still unclear if there is cell-to-cell variability in origin use and replication timing, and how this may link to genome instability, which could be addressed using the SPC-PTA workflow in parasites at different stages of the cell cycle. Additionally, the SPC-PTA workflow could be applied to capture rare recombination events that occur in the insect vector – events that contribute to *Leishmania* genetic diversity but are hard to catch with bulk methods. It has been demonstrated that *Leishmania* undergo a cryptic sexual recombination following meiotic patterns of genome inheritance^71–73^. However, gamete-like cells were never identified. Our tool could be applied to identify cells harboring whole genome loss-of-heterozygosity signatures, which would be a strong indicative of meiosis.

Another potential application is to combine the SPC-PTA workflow with loss-of-function assays to investigate key molecular factors involved in genome instability and DNA replication. For instance, small interfering RNA (siRNA) perturbations were combined with single-cell DNA sequencing to reveal a central role of Rap1 interacting factor 1 (RIF1) in coordinating replication timing in mouse embryos^74^. Although some *Leishmania* species lack a fully functional RNA interference pathway ^75^, loss-of-function assays can be achieved with CRISPR-Cas9-mediated gene knock-out^76^. More recently, genome-wide perturbation assays were made possible using CRISPR-Cas base editors opening the door to systematic, high-throughput functional screens in *Leishmania*^77,78^. Coupling these perturbation strategies with SPC-PTA could therefore enable direct testing of how some genes may shape replication dynamics, chromosome segregation, and genome instability at single-cell resolution.

Lastly, the use of semipermeable capsules itself is also a key innovation and open the possibility of other applications. For instance, as we demonstrated, parasites remain alive after SPC encapsulation, and the semi permeable nature of SPCs allow exchange of medium nutrients and cell metabolites, allowing cultivation of parasites inside SPCs^79^. This could be used to grow parasite clones trapped in SPCs, which would prevent cross-clone competition, allowing the recovery of slow growing clones which would be otherwise outcompeted by faster replicating clones. This is particularly important, e.g., for the study of quiescence, an emerging field of interest in *Leishmania* and other pathogens due to its adaptive role^80–83^, as it would allow to cultivate and separate parasites with a low replicative phenotype.

In conclusion, our high-throughput single-cell genome sequencing method provides an unprecedented view of *Leishmania* genome variation at cellular resolution, confirming previous observations of mosaic aneuploidy as well as revealing novel insights on the extend of CNVs and nucleotide sequence diversity within parasite populations. By accurately capturing structural variations alongside sequence polymorphisms, this approach offers a promising tool to link genetic variation to phenotypic outcomes (such as drug resistance or virulence) in protozoan parasites. More broadly, the SPC-PTA workflow has the potential applicability to other pathogens and cell systems that exhibit genome instability suggesting a wide impact in fields ranging from microbiology to oncology. In summary, the SPC–PTA workflow fills a critical gap in our experimental arsenal to reveal the interplay between genome instability and the hidden diversity that underlies parasite adaptation, drug resistance, and survival in changing environments.

## DATA AVAILABILITY

Raw sequencing data have been deposited in BioProject (NCBI) with accession number PRJNA1309081. Custom scripts used in this paper are available at Zenodo (https://doi.org/10.5281/zenodo.22876495).

## ACKNOWLEDGEMENTS

The computational resources and services used in this work were provided by the VSC (Flemish Supercomputer Center), funded by the Research Foundation - Flanders (FWO) and the Flemish Government.

## AUTHOR CONTRIBUTIONS

**G.H.N.**: conceptualization, data curation, formal analysis, funding acquisition, investigation, methodology, software, validation, visualization, writing – original draft, writing – review & editing. **P.M.**: Data curation, formal analysis, investigation, software, validation, visualization, writing – original draft, writing – review & editing. **J.C.D.**: conceptualization, funding acquisition, resources, writing – review & editing. **M.A.D.**: conceptualization, funding acquisition, methodology, project administration, resources, supervision, writing – review & editing.

