## Supplementary Material for "A novel high-throughput single-cell DNA sequencing method reveals hidden genomic heterogeneity in the unicellular eukaryote *Leishmania*"

By

Gabriel H. Negreira[1], Pieter Monsieurs[1], Jean-Claude Dujardin[2], Malgorzata A. Domagalska[1,*]

[1] Experimental Parasitology Unit, Institute of Tropical Medicine Antwerp, Antwerp, Belgium.

[2] Molecular Parasitology Unit, Institute of Tropical Medicine Antwerp, Antwerp, Belgium.

**SUPPLEMENTARY MATERIALS AND METHODS**

**Preliminary test on lysis conditions**

For preliminary tests for lysis efficiency, parasites were harvested by centrifugation (1,000 × g, 5 min) and washed three times with PBS. Aliquots of $1.5 \times 10^6$ cells were resuspended in either ice-cold PBS or fixed by addition of 4.5 mL ice-cold methanol with gentle mixing, followed by incubation at –20 °C for 30 min. Cells were subsequently washed three times in ice-cold PBS and resuspended in 250 µL PBS. Then, each cell pool (fixed vs non-fixed) was distributed into 4 conditions to compare lysis efficiency:

1. standard KOH lysis (0.8 M KOH, 20 mM EDTA, 200 mM DTT, 25 °C, 15 min).
2. extended KOH lysis (0.8 M KOH, 20 mM EDTA, 200 mM DTT, 25 °C, 60 min).
3. heated KOH lysis (0.2 M KOH, 20 mM EDTA, 200 mM DTT, 99 °C, 10 min).
4. No lysis (negative control).

After incubation under agitation at 100 rpm, samples were centrifuged (10,000 × g, 1 min) and had half of their supernatant transferred to a new tube. Lysis efficiency was estimated by quantifying the amount of free-floating DNA in the supernatant with qPCR using the SensiMix SYBR No-ROX kit (Bioline). Primers targeting the *Leishmania* conserved sequence region were used: CS-for (5'-GTCTTGGCGGTTCAGTTCG-3') and CS-rev (5'-GACATTGTGGTTCGTCTGCTC-3'). A standard curve was generated from serial dilutions of genomic DNA from *L. donovani* BPK081/0 cl8 (range: 1 ng to 1 fg). Reactions were run on a LightCycler 480 (Roche) with the following program: 95 °C for 10 min, then 45 cycles of 95 °C for 30 s, 60 °C for 15 s, 72 °C for 15 s, followed by a melting curve. DNA concentration in supernatants was estimated by fitting Ct values to the standard curve.

**Preliminary evaluation of PTA as WGA method**

Preliminary tests for PTA efficiency were done using genomic DNA from *Leishmania donovani* BPK081 strain as test input. Human genomic DNA supplied with the ResolveDNA™ kit (BioSkryb Biosciences) served as control. DNA was serially diluted in EB buffer (Qiagen) to obtain working concentrations of 10 pg and 0.1 pg, corresponding to approximately 100 and 1 *Leishmania* genomes, respectively. Negative controls contained EB buffer only. Whole-genome amplification was performed using the ResolveDNA™ kit (BioSkryb Biosciences) according to the manufacturer’s protocol. Briefly, 1 µl of diluted DNA (10 pg or 0.1 pg) or control was combined with 2 µl Cell Buffer and subjected to PTA amplification in a thermocycler using the following cycling conditions: 30 °C for 10h, and 65 °C for 3 min, followed by hold at 4 °C. Amplified DNA was purified using magnetic bead cleanup and eluted in 40 µl. DNA yield was measured with the Qubit dsDNA HS assay (Thermo Fisher Scientific). Reaction efficiency was expressed as fold amplification relative to input DNA mass. DNA fragment size distribution was assessed using the TapeStation 4200 with D5000 HS ScreenTape (Agilent Technologies).

### SUPPLEMENTARY RESULTS

#### Preliminary test on lysis protocol and WGA

For the test, we performed a preliminary experiment using bulk samples to assess lysis efficiency. This was done by submitting parasites to either the standard lysis workflow or to one of the modifications mentioned above, and by estimating the concentration of free-floating DNA in the supernatant after lysis with qPCR (supplementary figure 1). In all conditions, methanol fixation led to a reduction in free-floating DNA, indicating a potential negative effect on lysis and/or on DNA availability. In addition, in all conditions free floating DNA was 20-40 times more present compared to the non-lysed control, suggesting successful lysis. Although the 1h incubation seemed to have led to the highest amount of DNA in the supernatant, this observation would need more replicates to be confirmed.

To test if PTA would work with *Leishmania* DNA, which is 100X smaller and has a higher GC content compared to human DNA, we first made a preliminary assay using 10 pg and 0.1 pg (representing the DNA of ~ 1 *Leishmania cell*) of both human and *Leishmania* DNA as input in the PTA protocol. By measuring the output DNA with QuBit, we saw an amplification level in the range between 139.200 to 146.400 times for the human DNA, and 40.400 to 51.200 times for *Leishmania* DNA, showing that PTA does work with *Leishmania* DNA albeit with a lower, but still robust efficiency (supplementary table 1 and supplementary figure 2). Lastly, we tested if the full PTA workflow, including lysis, would work with *Leishmania* parasites by performing a second experiment using FACS-sorted cells as input. Here, output DNA were in the range between 10.000 to 35.920 times higher compared to input, indicating that PTA successfully amplified the DNA of the cells (Supplementary Table 2 and Supplementary Figure 2).

### SUPPLEMENTARY FIGURES AND TABLES

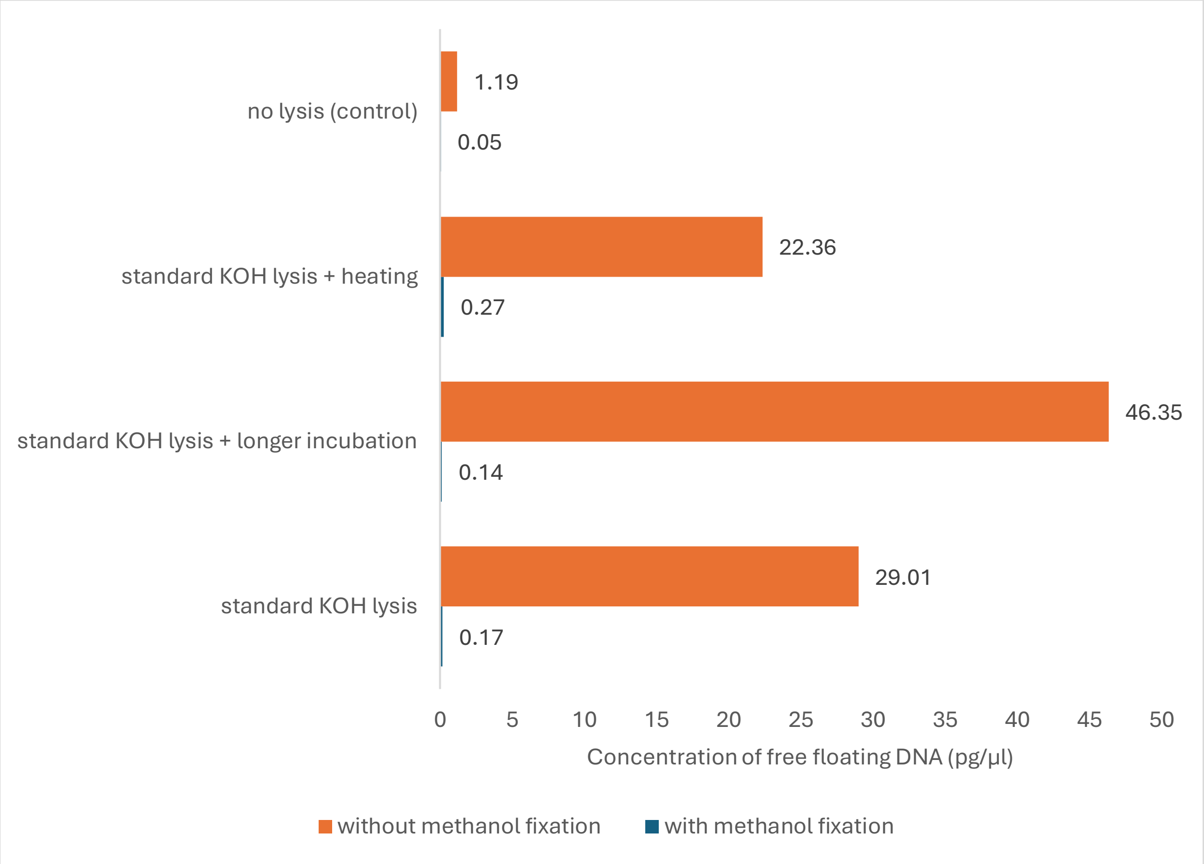

Supplementary Figure 1 – Evaluation of the efficiency of different lysis conditions. Bars indicate the concentration of free-floating DNA in the supernatant after promastigote pools were submitted to different variations of the lysis protocol of the Single Microbe Genome Barcoding kit. Concentrations were estimated with qPCR.

| Input | Concentration after WGA | Total mass after WGA | Output/input |
|---|---|---|---|
| 10 pg Human DNA | 36.6 ng/µl | 1464 ng | 146400 X |
| 0.1 pg Human DNA | 0.348 ng/µl | 13.92 ng | 139200 X |
| 10 pg *Leish* DNA | 10.1 ng/µl | 404 ng | 40400 X |
| 0.1 pg *Leish* DNA | 0.128 ng/µl | 5.12 ng | 51200 X |
| Negative Control | 0.0368 ng/µl | 1.472 ng | NA |

Supplementary Table 1 - Preliminary test of PTA with DNA samples. Output DNA concentration was estimated with QuBit.

| Input | Concentration after WGA | Total mass after WGA | Output/input |
|---|---|---|---|
| 100 cells - fresh | 8,62 ng/µl | 344,8 ng | 34480X |
| 10 cells - fresh | 0,25 ng/µl | 10 ng | 10000X |
| 100 cells -frozen | 4,96 ng/µl | 198,4 ng | 19840X |
| 10 cells frozen | 0,898 ng/µl | 35,92 ng | 35920X |
| Positive Control | 15,7 ng/µl | 628 ng | 62800X |
| Negative control | 0 ng/µl | 0 ng | NA |

Supplementary Table 2 -Preliminary test of PTA (including lysis protocol) with FACS sorted cells. For calculation of output/input ratios, Input mass was considered as 0.1 pg per cell. For frozen samples, cells were stored at -80˚C overnight before being submitted to PTA.

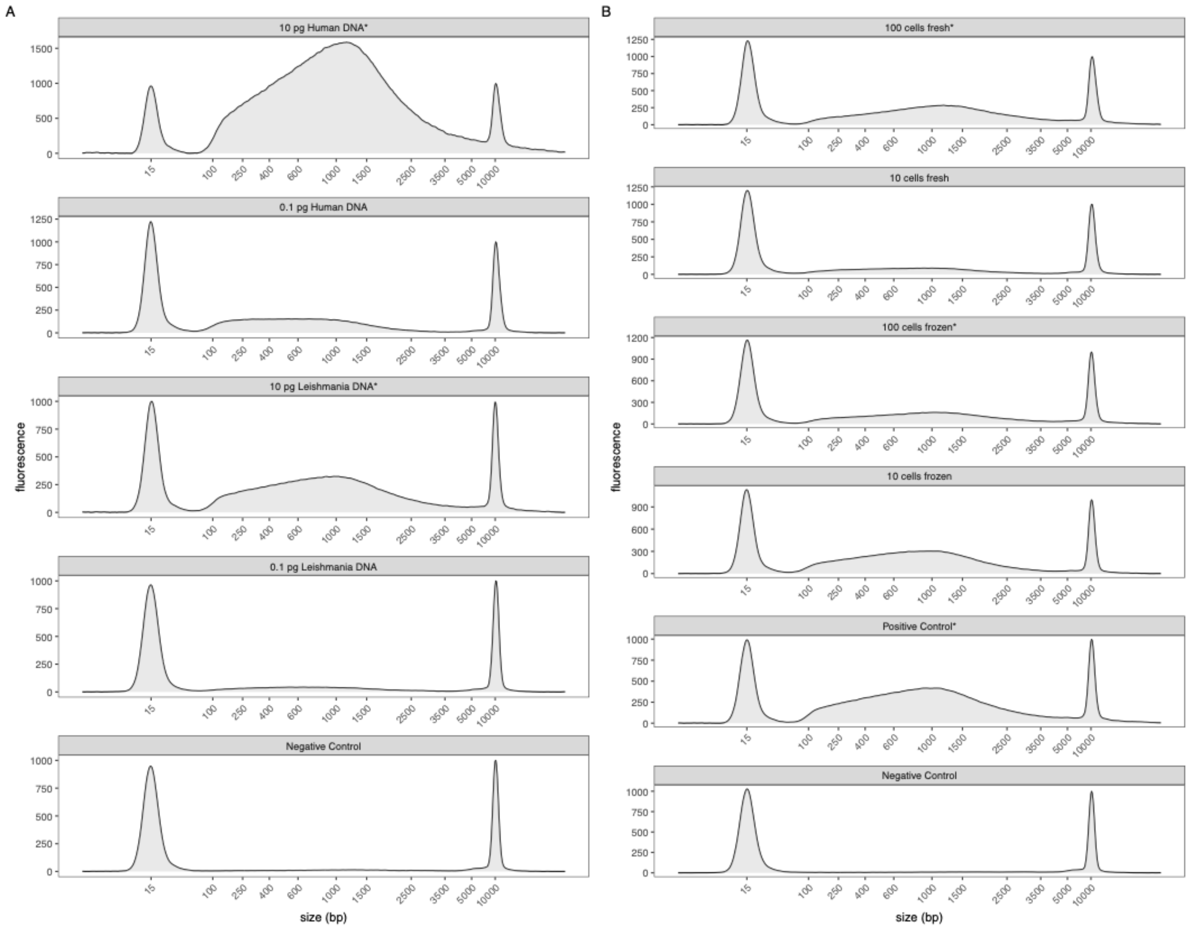

Supplementary Figure 2 - Preliminary test of PTA efficiency. A-B. Size distribution of amplicons of PTA products starting from DNA extraction samples (A) or FACS-isolated cells (B). * samples diluted 10X before loading. Positive control = 100 pg Leishmania DNA. Negative control = EB buffer.

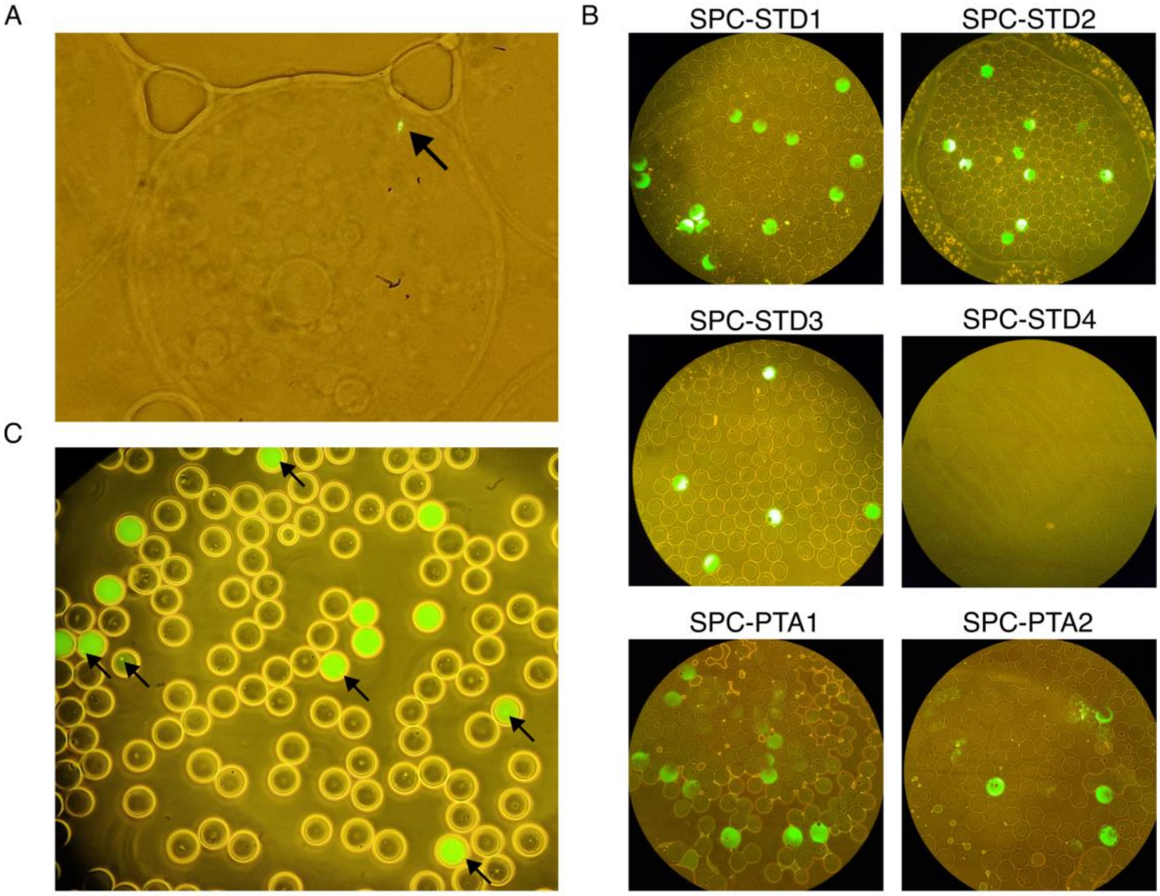

Supplementary Figure 3 – Fluorescence microscopy visualization of SPCs stained with SYTO5. A. Live *Leishmania* cell encapsulated in a SPC. B. Detection of amplified DNA after WGA with SYTO5. C. Presence of partially lysed cells after WGA on sample SPC-STD1.

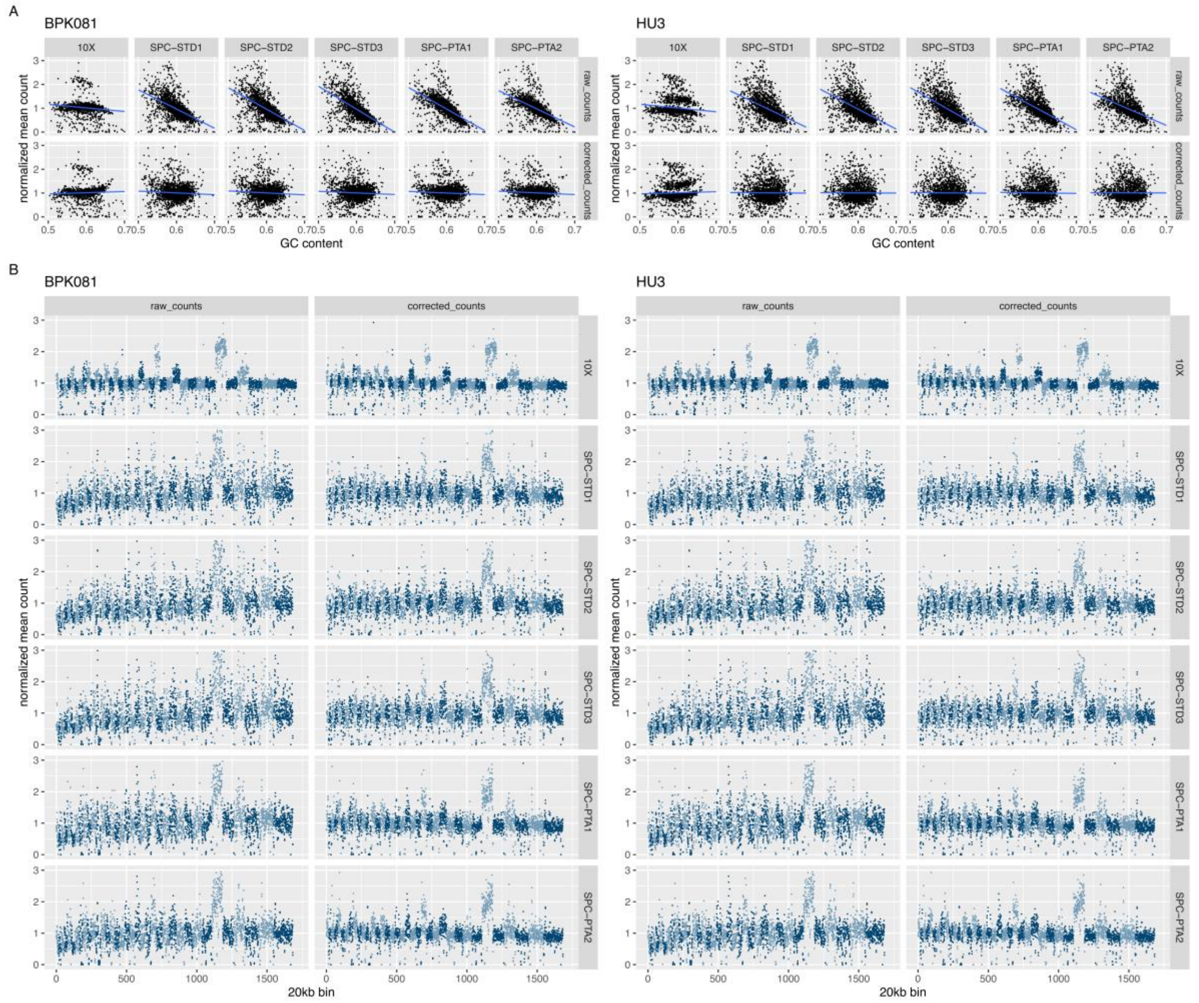

Supplementary Figure 4 - Effect of GC content on coverage. A. Correlation between the GC content (x-axis) and cell-normalized read count (y-axis) of 20 kb bins in BPK081(left) and HU3 (right) cells across all samples (columns). Top rows depict the raw counts, while bottom rows display the counts after GC bias correction. B. Coverage profile of combined reads from BPK081 (left) and HU3 (right) cells in each sample (rows) before and after GC-bias correction. Dots represent the combined cell-normalized read count in each 20 kb across the genome. Alternating blue colors were used to highlight chromosomes.

A

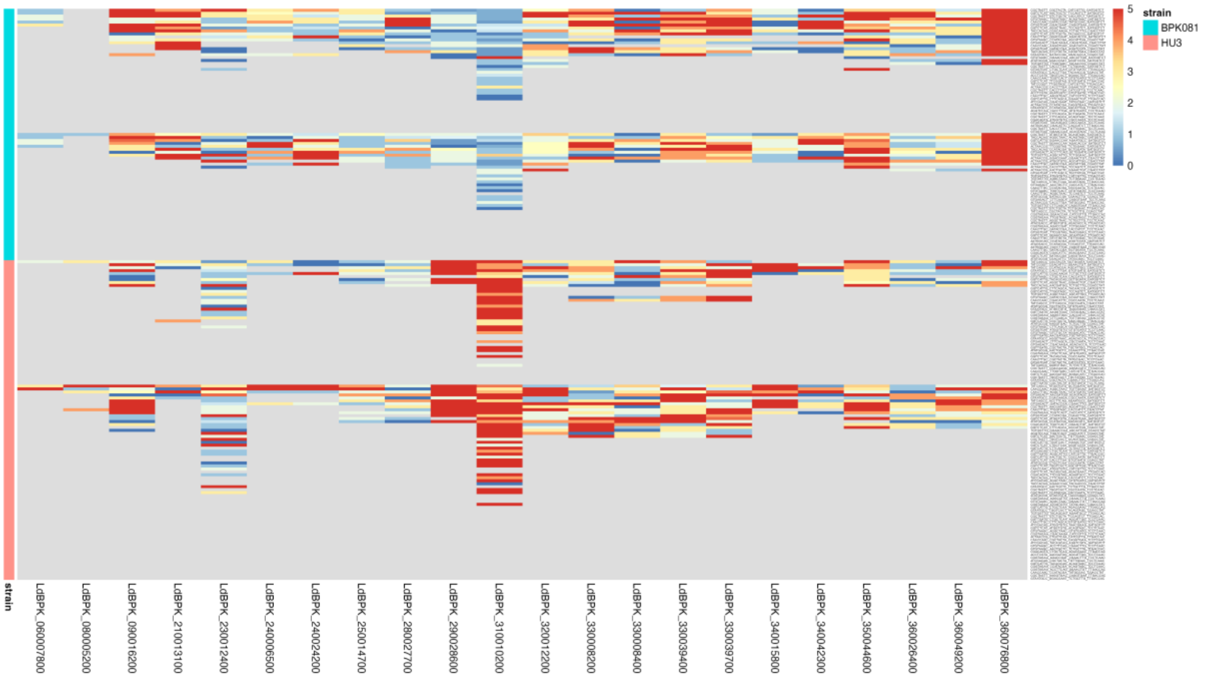

B

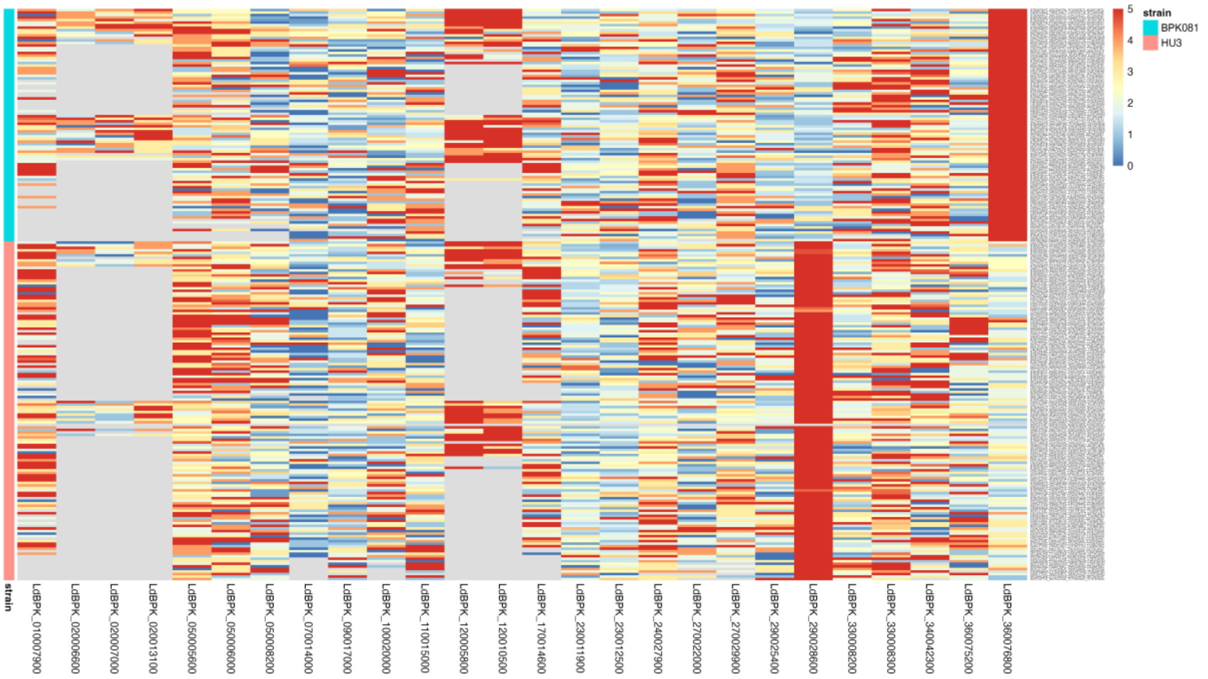

Supplementary Figure 5 – Detection of gene-level CNVs in the SPC-STD2 (A) and SPC-PTA2 (B) datasets. Colors in the heatmap indicate the estimated haploid copy number for each gene. Grey indicates missing values. Haploid copy number was estimated as the median read count in the annotated gene divided by the mean read count of the chromosome. Only genes with haploid copy number > 1.5 are shown.

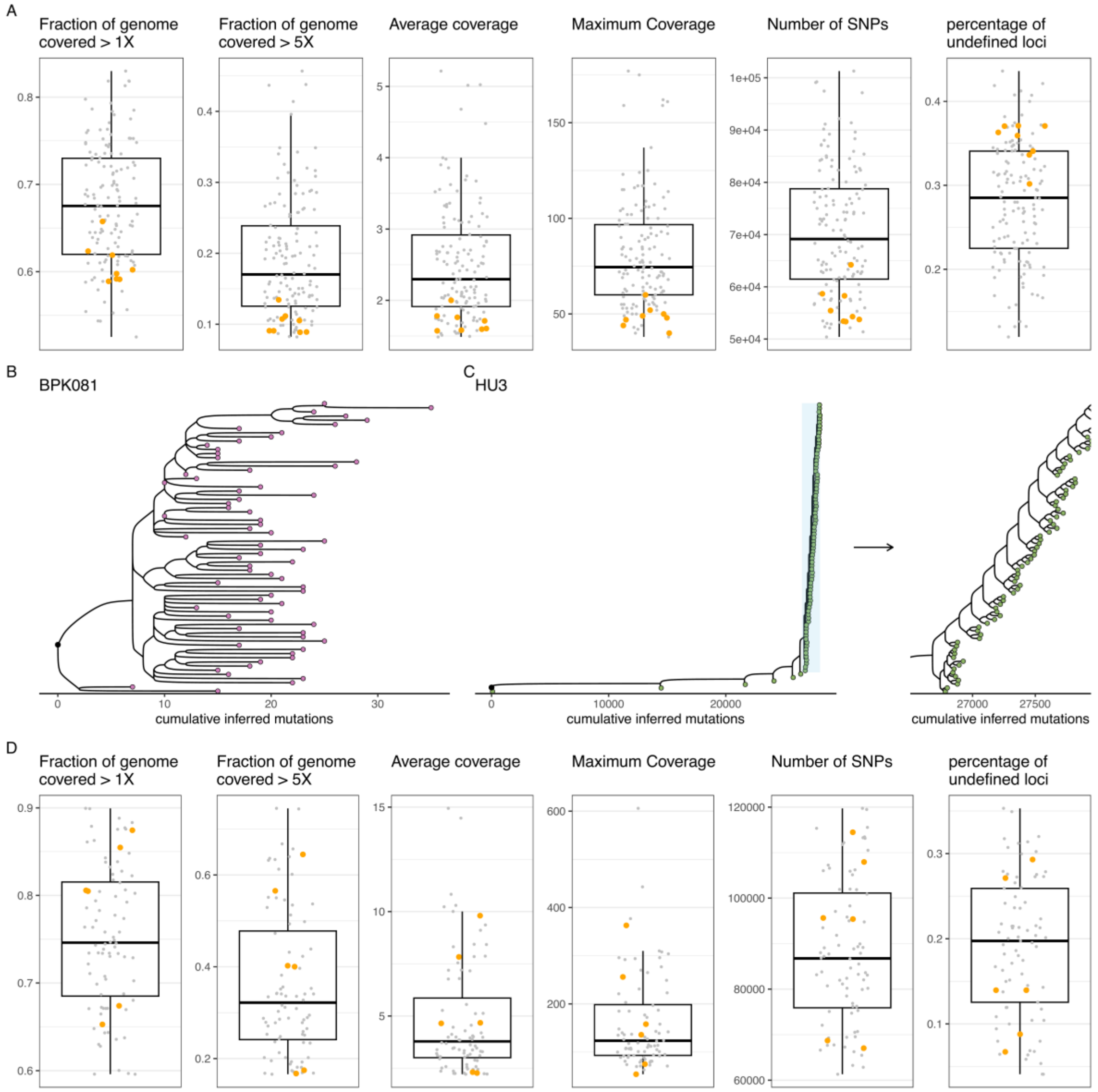

Supplementary Figure 6 – QC of diverging cells in HU3 phylogenetic trees. A. position of the 8 diverging cells (yellow dots) of the phylogenetic tree of HU3 from SPC-PTA2 in different coverage metrics compared to the other cells (grey dots). B-C phylogenetic tree of the BPK081 (B) and HU3 (C) cells in sample SPC-PTA1, constructed with scistree2. D. QC of the HU3 cells in SPC-PTA1. Yellow dots highlight the 6 diverging cells seen in the phylogenetic tree (figure C).
